# Spatial-transcriptomics reveals unique defining molecular features of fluorescence-sorted 5-aminolevulinic acid+ infiltrative tumor cells associated with glioblastoma recurrence and poor survival

**DOI:** 10.1101/2021.05.27.445977

**Authors:** Geoffroy Andrieux, Tonmoy Das, Michaela Griffin, Stuart J. Smith, Ruman Rahman, Sajib Chakraborty

## Abstract

Spatiotemporal-heterogeneity of glioblastoma (GBM) originating from the genomic and transcriptional variation in spatially distinct intra-tumor sites, may contribute to subtype switching in GBM prior to and upon recurrence. Fluorescence-guided neurosurgical resection utilizing 5-aminolevulinic acid (5ALA) has enabled the isolation of infiltrative margin tumor cells (5ALA+ cells) from a background of non-neoplastic cells. We have explored the spatial-transcriptomic (ST) landscape to interrogate molecular signatures unique to infiltrating 5ALA+ cells in comparison to GBM core, rim, and invasive margin non-neoplastic cells. ST analysis reveals that GBM molecular subtype plasticity is not restricted to recurrence, but manifests regionally in a cell-type-specific manner. Whilst GBM core and rim are highly enriched with Classical and Proneural subtypes, the unique enrichment of the Mesenchymal subtype (MES) in 5ALA+ cells supports the hypothesis that MES 5ALA+ cells may drive GBM recurrence. Upregulation of the wound response pathway in 5ALA+ cells signifies the possibility of hijacking the wound healing pathway of neural cells to promote tumor growth. Exon-intron split analysis revealed an upregulation of exonic counts for MES and wound-response genes in 5ALA+ cells, implying that these genes are under active post-transcriptional control. Network analysis suggests that wound response genes, including chemokine *CCL2* that recruits regulatory T-cells and monocytic myeloid-derived suppressor cells, are controlled by an *IRF8*-mediated transcriptional program in 5ALA+ cells. A higher stemness signature both in 5ALA+ cells and non-neoplastic cells of the invasive margin emphasizes the role of this microenvironment in stemness acquisition and defines 5ALA+ cells as a rare sub-population of GBM stem cells. Finally, we establish a link between the unique molecular signatures of 5ALA+ cells and poor survival and GBM recurrence. Characterization of the 5ALA+ infiltrative sub-population offers an opportunity to develop more effective GBM treatments and urges focus away from the GBM proliferative core, upon which failed targeted therapies have been predicated.

## Introduction

Isocitrate dehydrogenase [IDH]-wild-type Glioblastoma (GBM) is a highly aggressive and heterogeneous tumor with poor survival outcome. Despite radical multimodal treatment of aggressive surgery, radiation therapy, and chemotherapy with temozolomide, the poor treatment outcome of GBM patients has remained stagnant with a median survival of 14.6 months from diagnosis ^1^. One of the key reasons of poor treatment response is the invasiveness of GBM deep into the neighboring brain parenchyma, which renders complete surgical resection impossible and efficacious brain penetration of chemotherapeutics a considerable challenge ^2^. Furthermore, the ineffectiveness of therapeutic agents may arise from the plasticity of GBM cells which manifests intra- and inter-tumor heterogeneity; indeed, failed molecular targeted therapeutics have historically been focused on the GBM proliferative genotype ^3^. Heterogeneity in GBM has now been well established and contributes to the differential expression of subclonal genes owing to distinct molecular events developing during the spatiotemporal evolutionary lifespan of GBM cells ^4^. In search of the potential origins of phenotypic diversity and plasticity of GBM cells, the recent body of work indicates the existence of a rare GBM stem cell (GSCs) sub-population with self-renewing capacity ^5, 6^. Major characteristics of the GSCs include hijacking the normal neural stem cell developmental programs to promote and maintain tumor growth, and acquisition of mechanisms to resist chemotherapy ^7^. A recent single-cell RNA sequencing (scRNA-seq) study revealed a high inter- and intra-GSC transcriptional heterogeneity characterized by a transcriptional gradient composed of distinct gene signatures of two cellular states - „normal neural development‟ and „inflammatory wound response‟ and concluded that a transcriptional program resembling a neural wound response in GSC may functionally contribute to GBM initiation ^8^. To gain insight into the functional developmental and metabolic programs of GBM cells, Garofano *et al.* integrated scRNA-seq and bulk transcriptomics data by using a computational platform - single-cell biological pathway deconvolution (scBiPaD), and showed that the distribution of GBM cells along neurodevelopmental and metabolic axes, could facilitate their classification as proliferative/progenitor, neuronal, mitochondrial (MTC) and glycolytic/plurimetabolic (GMP) subtypes ^9^.

Based on transcriptional attributes, GBM was originally classified into four subtypes: Classical (CL), Neural (NE), Proneural (PN), and Mesenchymal (MES) subtypes ^10^. A more recent study based on GBM transcriptomics analysis, excluding nonmalignant cell types, confirmed three subtypes of GBM – CL, PN, and MES ^11^. Emerging evidence has revealed the plasticity of GBM subtypes suggesting that most patients exhibit varying subtypes within the same tumor ^4^. Moreover, longitudinal studies demonstrated the temporal plasticity of GBM subtypes by uncovering the subtype switch in GBM patients upon recurrence ^11, 12^. Recently, Minata *et al.* showed that in response to the radiation-induced proinflammatory microenvironment, GBM cells at the edge of the tumor acquire a MES subtype defined by the expression of CD109 ^13^. This differential expression of CD109, observed between paired primary and recurrent patient-cohorts, is assumed to be associated with MES transition upon GBM recurrence ^13^.

Apart from temporal plasticity, GBM exhibits spatial heterogeneity and a growing body of evidence suggests that a collection of tissue from a single site fails to capture the spatial heterogeneity and transcriptional dynamics of GBM cells ^14^. Tissues isolated from the resected tumor margin where GBM penetrates into the normal brain parenchyma, harbors distinct genomic and transcriptomic profiles in contrast to tissue removed from tumor core and enhancing rim regions ^14^. Recurrence of GBM is initiated at the invasive margin within 2cm of the resected region after surgery ^15^. Therefore, genomic and transcriptomic profiles of the invasive margin have the unique potential to identify molecular candidates which can be targeted to impair the recurrence of GBM. Nevertheless, the highly heterogeneous cellularity of the tissues taken from the invasive margin, including infiltrated immune and normal neural cells, pose a substantial challenge to filter the tumor-specific genomic and transcriptomic profiles from an overwhelming background of non-neoplastic cells. A viable solution emerged from the use of 5-aminolevulinic acid (5ALA) during GBM neurosurgery, reported to increase the percentages of complete resection from 36% to 65% ^16^. 5ALA - a porphyrin - is metabolized to a fluorescent metabolite protoporphyrin IX (PpIX) by cells in which the heme biosynthetic pathway is activated, such as GBM cells but not non-neoplastic cells ^14^. GBM cells exhibit a marked accumulation of PpIX due to their inability to incorporate iron into the protoporphyrin core because of the absence of ferrochelatase enzyme activity. The accumulated PpIX in GBM cells show a maximum excitement by blue light at 400-410 nm with an emission light peak at 635-704 nm exhibiting a pink fluorescence corresponding to regions of viable tumor cells ^14^. The necrotic tumor core does not emit fluorescence owing to their deficiency of active heme metabolism; in contrast to the invasive tumor margin which fluoresces brightly and eventually fades with the decreasing number of tumor cells in the periphery ^17^. We have shown that invasive margins from primary tissue harboring the infiltrative 5ALA+ tumor cells can be purified from background non-neoplastic cells through fluorescence-activated cell sorting (FACS) ^14^.

Intra-tumor surgical sampling, aided by FACS-isolation of 5ALA+ and 5ALA-subpopulations from invasive margin tissue, can serve as an ideal resource to untangle the intricacy of cellular and transcriptional heterogeneity within GBM. We have demonstrated that analysis of spatial transcriptomics within this sampling framework can identify non-canonical molecular factors associated with GBM infiltration, such as *SERPINE1* ^14^. However, deep profiling of transcriptomic landscapes of spatially distinct tumor core, rim, invasive margin, and 5ALA sorted cells, is required to elucidate the spatial transcriptional heterogeneity and unique molecular signatures of 5ALA+ cells.

Since 5ALA+ cells represent a rare tumor subpopulation within the invasive margin and importantly, are in closest proximity to residual disease remaining post-surgery, it is pertinent to explore the molecular subtypes, cellular states, and immune-related pathways in 5ALA+ cells and how this differs from 5ALA-non-neoplastic cells and distinct intratumor regions of GBM. Elucidating molecular signatures of emerging cellular (developmental and inflammatory wound response) and metabolic (GPM and MTC) states in 5ALA+ cells, is crucial to better understand the role of these infiltrative cells in tumor promotion and maintenance. Moreover, interrogating this rare subpopulation for stem-cell like features can also shed light on survival outcome and recurrence of GBM.

To address these questions, we performed spatial-transcriptomics (ST) analyses of unsorted tumor core, rim, and invasive margin tissue, and FACS-isolated 5ALA+ and 5ALA- cells across 10 GBM patients. We interrogated the transcriptomic landscape for molecular, cellular, metabolic and stemness gene signatures, to test the hypothesis that the 5ALA+ subpopulation(s) is defined by a unique cellular state. To understand the transcriptional programs of 5ALA+ cells, we employed exon-intron split analysis (EISA)^18^ to distinguish between transcriptional and post-transcriptional control mechanisms and ARACNE algorithm to construct the transcriptional network governing unique gene signatures of 5ALA+ cells. Lastly, we explored the association of 5ALA+ cells with survival outcome and recurrence of GBM.

## Methods and Materials

### Tissue sample collection, processing, and FACS analysis

Tissue samples representing spatially distinct regions (tumor core, rim, invasive margin) of tumor samples from 10 GBM patients were retrieved that were previously collected as part of an earlier study ^14^. The consent and ethical approval were obtained from the Local Regional Ethics Committee in Nottingham (East Midlands Ethics Approval Reference: 11/EM/0076; 5th May 2011). Briefly, 5ALA (20 mg/kg dose) was administered orally to patients 2–4 h prior to craniotomy and visualization of 5ALA induced fluorescence. Aided by image guidance multi-region tissue samples were collected. For instance, samples were taken from non-fluorescent or minimally fluorescent regions representing tumor core, whilst samples from the viable fluorescent region corresponded to tumor rim. The furthest region of 5ALA-induced fluorescence beyond the bulk tumor, where the tumor penetrated adjacent healthy parenchyma, corresponded to the invasive margin. Histological diagnosis and formal postoperative diagnosis (including IDH mutations, ATRX mutation, and MGMT methylation status) were investigated.

Cells were dissociated from the invasive margin and subjected to Fluorescence-Activated Cell Sorting (FACS) based on 5ALA immunofluorescence as described by us previosuly ^14^. Briefly, the cells were sorted using an excitation spectrum at 405 nm and an emission spectrum at 605–625 nm. The purity of the cells was ensured, by sorting the cells twice consecutively. As controls, U251 GBM cells incubated for 2 h with and without 5ALA were used for FACS gating. The positive and negative sorted cells were subsequently centrifuged at 800 rpm (180 × g) for 5 min before subjected to snap freezing.

### Immunohistochemistry

Tissues from spatially distinct regions of GBM were collected and immunohistochemistry performed as described by us previously ^2^. Samples were obtained from tumor core (superficial and anterior medial), tumor rim (deep edge), and invasive margin. Briefly, after the removal of paraffin wax, samples were treated with sodium citrate buffer (pH 6) for 40 minutes at 90 °C and washed with phosphate buffer solution (PBS) for 2 minutes. Then, 200 μL of peroxidase blocking solution was applied to cover the specimen for 5 minutes followed by washing with PBS. After the slides were dried, Ki-67 antibody (DAKO) and CD31 (DAKO) were applied with 1:50 dilution and incubated for 1 hour at room temperature. Sections were washed with PBS before the addition of the secondary antibody (DAKO) and incubated at 37°C for 30 minutes. Finally, the substrate-chromogen solution (DAB) was applied to cover the specimen, incubated for 5 minutes, and rinsed gently with distilled water. The Olympus BX41 light microscope was used to visualize and capture the images of the different GBM regions.

### RNA isolation from spatially distinct regions of GBM

Dissociation for tumor core, rim, and the invasive margin was performed by a previously described by us ^19^ Total RNA extraction followed by quality control analysis was performed as described by Smith *et al.* ^14^. Briefly, libraries were prepared using the NEBNext Poly(A) mRNA Magnetic Isolation Module (NEB: E7490), the NEBNext Ultra Directional Library Kit for Illumina (NEB: E7420) and the NEBNext Multiplex Oligos for Illumina (Index Primers Set 1) (NEB: E7335L). Samples with total RNA concentration of >10 ng/µl (0.5 µg total amount) were used for library preparation. To ensure library quality, adequate concentrations were obtained from each sample, followed by 14 cycles of amplification during the PCR-based library enrichment step.

Finished libraries were quantified using the Qubit dsDNA HS kit (Invitrogen: Q32854). Library concentrations as well as fragment size distributions were also analyzed by employing the Agilent Bioanalyzer High Sensitivity DNA Kit (Agilent: 5067-4626).

Libraries were normalized to 2 nM and pooled in equimolar amounts. The Kapa Library Quantification Kit (KAPA Biosystems: KK4824) was used for precise quantification of the library pool. The library pool was denatured and diluted to 1.6 pM, spiked with 1 % PhiX (1.8 pM) and sequenced on the Illumina NextSeq 500, using the NextSeq 500/550 High Output v2 Kit (150 cycles) (Illumina: FC-404-2005), to generate a minimum of 70 million pairs of 75-bp paired-end reads per sample. Raw RNA-seq data has been deposited at ArrayExpress with accession number E-MTAB-8743.

### RNA-seq analysis

We obtained RNA-seq raw data (FASTQ files) from spatially distinct unsorted regions (tumor core, tumor rim, and invasive margin) and 5ALA sorted cells from invasive margin across 10 GBM patients. RNA-seq raw data from spatially distinct regions were processed by using Bioconductor package QuasR (version 1.30.0) ^20^. For primary alignment, we used the reference genome hg19 for human in this work. The QuasR package employs the required tools to obtain expression tables from the raw RNA-seq reads and includes the aligners Rhisat2 ^21^ and SpliceMap ^22^. We performed the alignment by using the following command:

„qAlign(“sampleFile.txt”,“BSgenome.Hsapiens.UCSC.hg19”,splicedAlignment=TRUE, aligner=“Rhisat2”)‟.

We then measured the count of each gene within any annotated exonic region using the function qCount. The count data obtained from the QuasR package was then converted to transcripts per million (TMP) followed by log transformation (Log_2_) of TPM+1 values. Initial quality control analysis was performed on the normalized RNA-seq counts including a comparison of differential expression across the samples. Genes with valid count values were additionally compared among the samples.

### Gene-set enrichment analysis (GSEA) Hallmark enrichment

Enrichment analyses of the hallmark gene sets representing the well-characterized biological processes related to cancer were carried out by a gene set enrichment analysis (GSEA) algorithm ^23^. Briefly, the hallmark gene sets were selected from MSigDB gene-set collections ^24^, and enrichment analysis was conducted amongst the different regions (tumor core, tumor rim, and invasive margin) and 5ALA sorted cells (5ALA+ and 5ALA- cells) using GSEA. The ranked list of genes obtained from GSEA was further processed by Fast Gene Set Enrichment Analysis (fgsea) R-package ^25^. Normalized enrichment score (NES), p-value, and adjusted p-values (calculated with a standard Benjamini-Hochberg - BH procedure) were retrieved for each of the hallmarks that were enriched in different regions and cell populations. The hallmarks with higher NES values and adjusted p-value < 0.05 were considered as enriched for a specific GBM region.

For further analysis of the enriched pathways, the leading edge genes representing the subset of genes contributing significantly to the enrichment signal of a given gene set ^23^ in a specific GBM region, were identified and subjected to hierarchical clustering analysis. We employed a combined ComplexHeatmap::pheatmap() function that uses two R-packages, „pheatmap‟ and „ComplexHeatmap‟, for the clustering purpose where the column and row clustering were performed by using the Euclidean distance method.

### Neural cell-type gene-signature enrichment

To characterize the different GBM regions, transcriptome-based neural cell-type signatures described by Cahoy *et al.* were retrieved ^26^ (Supplementary Table S1). In brief, Cahoy *et al.* employed Affymetrix GeneChip Arrays to identify gene-signatures of different neural cell types including neurons, oligodendrocytes, astrocytes, and cultured astroglial cells ^26^. NES and adjusted p-values were calculated using GSEA and fgsea algorithms as described above.

### GBM subtype gene-signature enrichment

Gene signatures of each GBM molecular subtype were obtained from Verhaak *et al.* which described an efficient gene expression-based molecular classification of GBM samples into four molecular subtypes: Proneural (PN), Neural (NE), Classical (CL), and Mesenchymal (MES) ^10^ (Supplementary Table S1). The signature gene-set for each of the subtypes was retrieved from MSigDB. GSEA for the molecular subtypes was performed on the spatially distinct GBM regions. NES and adjusted p-values were calculated using the fgesa package as described previously.

### Developmental, inflammatory wound response, and metabolic gene signature enrichment

Signature genes for developmental and inflammatory wound healing/ injury response phenotypes were retrieved from Richards *et al.* ^8^ (Supplementary Table S1). In addition, two gene-sets representing two divergent metabolic phenotypes – mitochondrial (MTC), glycolytic/plurimetabolic (GPM) were retrieved from Garofano *et al.*^9^ (Supplementary Table S1). GSEA was performed to identify the enrichment of these diverse gene-sets in spatially distinct GBM regions.

### Stemness gene-signature enrichment

Diverse signature gene-sets representing stem cells ^27^, cancer stemness ^28^, embryonic stem cells (ES1 and ES2) ^29^, Human Embryonic Stem Cells (hESC) ^30^, induced pluripotent stem cells (iPSC) ^31^, Nonog/Sox2 induced stem cell gene-set ^29^, Myc induced ES gene-set ^32^ and Human Epithelial ASC ^33^ were retrieved (Supplementary Table S1). GSEA was performed to identify the enrichment of these diverse gene-sets in spatially distinct GBM regions as described earlier.

### Estimation of mRNA expression-based stemness index (mRNAsi) in spatially distinct regions of GBM and TCGA-GBM samples

Estimation of mRNA expression-based stemness index (mRNAsi) was performed by the method described by Malta *et al.* ^34^. Briefly, to calculate mRNAsi, a machine learning approach was used to develop a predictive model by employing one-class logistic regression (OCLR) as described previously by Sokolov *et al.* ^35^. The OCLR was based on the hESC and iPSC from the Progenitor Cell Biology Consortium (PCBC) dataset ^36, 37^. The mRNAsi score ranges between 0 and 1, where 0 indicates less stemness with a more differentiated tissue state and 1 represents more stemness with a less differentiated state. A gene expression matrix (samples in columns and genes in rows) was prepared and employed on R-package “TCGAbiolinks” using the function TCGAanalyze_Stemness() to generate a stemness score based on the spatial RNA-seq data from 10 patients.

### Construction of the transcriptional network

Firstly, the transcription factors in the leading gene sets representing the cellular and metabolic states including inflammatory pathway, TNFA signaling via NFKB pathway, MES subtype, inflammatory wound response, MTC, and GPM subtypes were manually curated by using the ENCODE database ^38^. We used the mutual information-based algorithm ARACNE (Algorithm for the Reconstruction of Accurate Cellular Networks) ^39^ to construct the regulatory network between transcription factors and target genes based on mRNA expression values of the leading edge genes. The previously described bootstrap algorithm ^40^ was used to assess statistical confidence. We inferred 1000 networks based on bootstrap datasets, setting the most stringent value for Data Processing Inequality (DPI = 0) tolerance. Finally, we estimated the significance of the edges by testing their probability against a null distribution obtained by random permutation of predicted edges. The consensus network conserves edges with a p-value below 10^-4^. Based on the ARACNE output we retrieved only the mutual information (MI) values for a given transcription factor and target genes for which the p-value was significant (<0.05), and constructed and visualized the transcription factor – target genes network by using Cytoscape ^41^.

### Survival analysis

Survival analysis based on TCGA mRNA Log_2_ expression data (TPM+1) of the gene signatures was performed by using GEPIA2 ^42^. Briefly, the survival curves were generated using Kaplan–Meier analysis for overall (OS) and disease-free survival by using a median survival cutoff. The hazard ratios were estimated by Cox proportional hazards model regression analysis ^43^. Analysis was performed at a 95% confidence interval. Survival analysis for each of the gene signatures representing inflammatory response, TNFA signaling, MES subtype, inflammatory would response, and metabolic subtypes (GPM and MTC) was performed separately. For classifying the high- and low-expression cohorts, the median expression threshold was set (Ccut-off high: upper 50% and cut-off low: lower 50%).

### Exon-intron split analysis (EISA)

Exon-intron split analysis (EISA) was employed as described by Gaidatzis *et al.* ^18^ to investigate the changes in pre-mRNA (introns) and mature-mRNA (exons) counts across different regions of GBM, that leads to the quantification of transcriptional and post-transcriptional control of gene expression. R-package “eisaR” was used for the EISA. Briefly, after mapping the transcripts to a unique position in the genome, we quantified counts of annotated exonic readsrepresenting mature mRNAs and the count of the reads that did not match any annotated exons (intronic). Normalization was performed for exons and introns separately by dividing each sample by the total number of reads and multiplying by the average library size. Based on these expression levels only the genes with reasonable counts (average log_2_ expression level of at least 5) were selected for downstream analysis. Genes with overlapping reads were discarded as it is difficult to assign intronic reads to the respective genes. A differential exonic and intronic change among different regions has been performed with EdgeR, as described by Gaidatzis *et al.* A p-value < 0.05 was considered significant.

GSEA of Hallmark, 5ALA+ cell-derived signatures, and transcription factors were conducted as described before, with genes ranked based on the log_2_ difference of exon or intron normalized intensity between two regions used as input.

### Recurrent *vs.* Primary GBM analysis

To compare the gene-signature enrichments in IDH wild-type recurrent and primary GBM tumors, RNA-seq data was retrieved from The Cancer Genome Atlas (TCGA) (N = 140, Primary = 128 and Recurrent = 12) (https://tcga-data.nci.nih.gov/tcga/), Chinese Glioma Genome Atlas (CCGA) (N= 190, Primary = 109 and Recurrent = 81) (http://www.cgga.org.cn/) ^44^ and The Glioma Longitudinal AnalySiS (GLASS) (N= 60, Primary = 30 and Recurrent = 30) ^45^ cohorts. Gene signatures comprised of the leading edge genes representing TNFA signaling, inflammatory response, MES subtype, inflammatory wound response, MTC subtype, GPM subtype, stemness and transcription factors that were upregulated in 5ALA+ cells, were used for enrichment. GSEA was performed to identify the enrichment of these diverse gene sets in spatially distinct GBM regions as described earlier. In order to investigate the correlation between the unique

5ALA+ gene-signatures and survival of GBM patients, a single sample GSEA ^46^ was performed to identify the enrichment of Mesenchymal and Wound Response pathways for each primary and recurrent tumor samples from TCGA, CCGA and GLASS cohorts. Normalized enrichment scores for each of the gene sets were accumulated to calculate the 5ALA+ gene-signature score for each primary and recurrent GBM patient. The Spearman correlation coefficient was calculated between the 5ALA+ gene-signature scores of GBM patients and survival data for primary and recurrent GBM separately.

## Results

### Correlation analysis reveals transcriptome compatibility of unsorted GBM tissues and FACS-isolated cells

Due to the distinct nature of the unsorted tissue and sorted cells (i.e. FACS process in the latter), we performed quality control analysis to evaluate the compatibility of the transcriptome between unsorted tissues - tumor core (Core), tumor rim (Rim), and invasive margin (Inv) and sorted cells (5ALA+ and 5ALA-). The analysis showed that the log_2_ normalized RNA-seq counts (TPM+1) were uniform and showed less variance across different patients in the unsorted regions. Inter-region variability did not supersede the intra-region patient variability in terms of mRNA-expression. However, a notable variation was observed between the 5ALA sorted cells and the unsorted tumor regions (Core, Rim, and Inv) (Supplementary Figure S1A). The underlying cause for these differences between the unsorted tissue regions and sorted cells could be attributable to both biological and technical factors. The unsorted regions represent heterogeneous cell populations within the tissue, whereas the sorted cells were isolated and subjected to FACS analysis. To evaluate the compatibility further, we calculated the person correlation coefficients on the normalized mRNA expression data amongst the different regions and 5ALA sorted cells. Analysis indicated that correlation values across different tumor regions and cells were uniform ranging from 0.88 to 1.00. When the expressions of the shared genes were compared between unsorted regions and sorted cells, a reasonable correlation was observed (Core *vs.* 5ALA-: 0.92, Rim *vs.* 5ALA-: 0.93, Inv *vs.* 5ALA-: 0.91, Core *vs.* 5ALA+: 0.89, Rim *vs.* 5ALA+: 0.89, and Inv *vs.* 5ALA+: 0.88) (Supplementary Figure S1B). A higher correlation was also observed between 5ALA- and 5ALA+ cells (R= 0.95) as predicted by a shared brain microenvironment. The highly correlated transcriptome of the unsorted tissue and sorted cells were indicative of their compatibility.

### Differential regulation of cancer-specific and metabolic pathways in spatially distinct GBM regions

Having established the compatibility of the ST datasets from unsorted tissues (Core, Rim and Inv) and 5ALA sorted cells (5ALA+ and 5ALA-), we performed a diverse array of computational analysis to identify unique defining molecular features of 5ALA+ cells relative to unsorted tissues and 5ALA- cells (Figure 1A). To determine the cancer-related and metabolic pathways which are associated with distinct regions of GBM (Core, Rim, Inv) and 5ALA sorted cells from the Inv margin, a GSEA was performed (Supplementary Table S2) for cancer-hallmarks. Tumor core exhibited the highest number of enriched pathways including Epithelial-mesenchymal transition and Hypoxia, which had the highest normalized enrichment scores and the lowest adjusted p-values (Supplementary Table S2), followed by TNFA signaling via NFKB, Interferon-alpha response, Inflammatory response, E2F targets, and IL-6/JAK/STAT3 signaling (Figure 1B). Akin to the Core, Epithelial-mesenchymal transition and Hypoxia were highly enriched in the Rim (Figure 1B and Supplementary Table S2). The Core was also enriched with pro-proliferative pathways such as mitotic-spindle, G2M checkpoint, mTOCR1 signaling and E2F targets, whereas Inv margin was devoid of pro-proliferative pathways (Figure 1B). To investigate the distribution of proliferative cells across spatially distinct GBM regions, Ki-67 immunohistochemistry (IHC) was performed, revealing a high number of proliferative cells in the Core superficial medial region (Figure 1C) followed by anterior medial (Figure 1D) and tumor-edge regions (Figure 1E). In contrast to the Core, the GBM Inv margin exhibited a lower number of proliferative cells (Figure 1F). The pro-proliferative pathways enriched in the Core, underlie the increased number of proliferative cells observed in this region compared to the Inv margin. The enrichment of glycolytic and angiogenesis pathways (Figure 1B and Supplementary Figure S2A and B) in addition to the absence of oxidative phosphorylation in the Core and Rim, supports a Warburg-like effect induced by the hypoxic conditions in these regions (Figure 1B and Supplementary Figure S2C and D). In contrast, Hypoxia was neither enriched in the unsorted Inv margin, nor 5ALA+ and 5ALA-sorted cells. Enrichment of oxidative phosphorylation in Inv margin further corroborated the evidence suggesting the GBM infiltrative margin represents a non-hypoxic microenvironment (Figure 1B). CD-31 IHC also reinforced this finding by showing that Core and Rim were highly vascularized (Figure 1G and H), whereas no vascularization was observed in Inv margin (Figure 1I). Interestingly, TNFA signaling via NFKB and Inflammatory response pathways were highly enriched in 5ALA+ cells (Figure 1B, Supplementary Figure S2E, and S2F and Supplementary Table S2).

**Figure 1:**
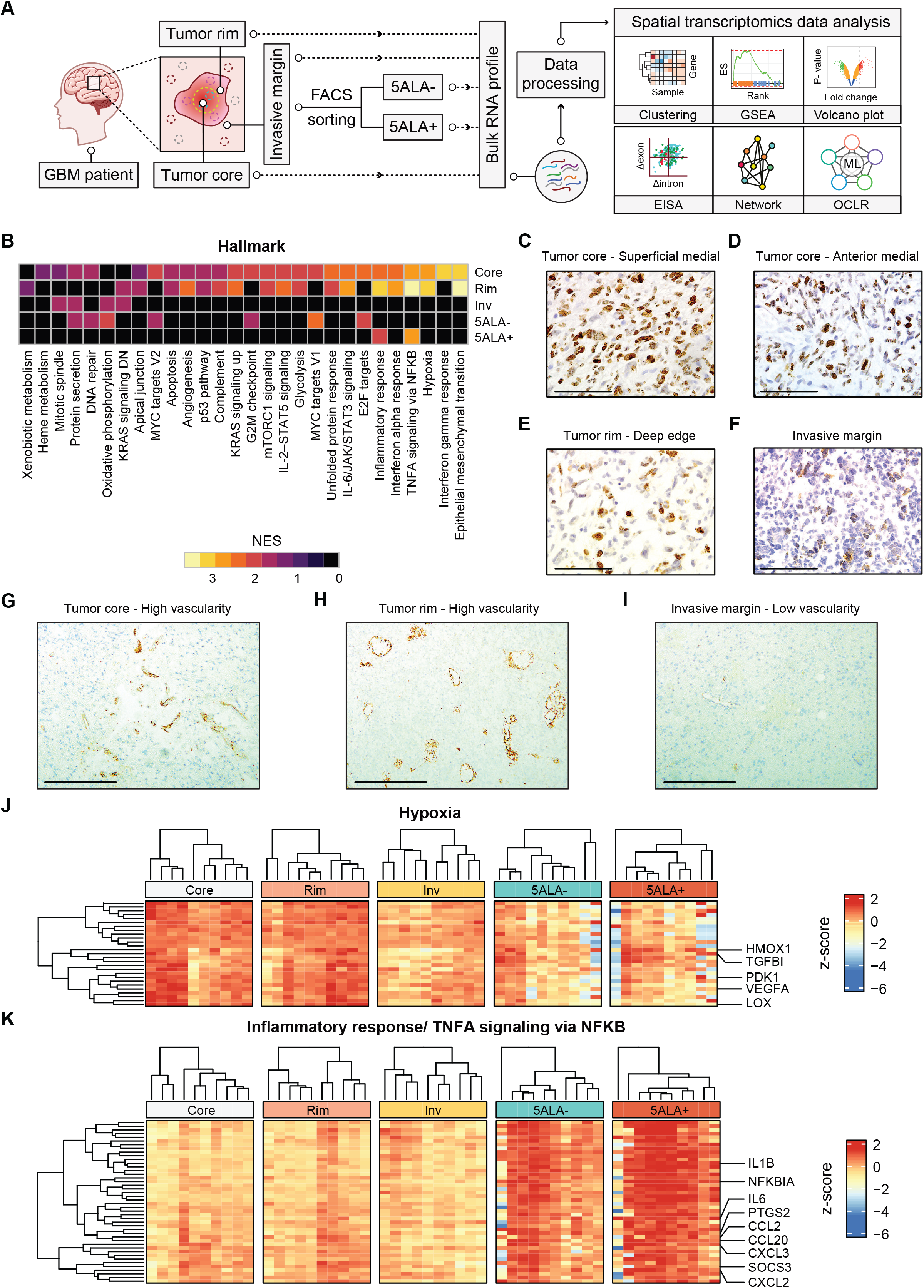
Hallmark gene set enrichment analysis (GSEA) in spatially distinct different GBM. Schematic figure delineating the steps for tissue collection from distinct GBM regions (tumor core, tumor rim and invasive margin) followed by 5ALA flouresence-based FACS isolation of invasive margin cells into 5ALA+ and 5ALA-subpopulations. The spatial-transcriptome from each unsorted region and sorted cells were interrogated by gene-set enrichment analysis (GSEA), hierarchical clustering, differential expression analysis, Exon-intron split analysis (EISA), network inference and one class logistic regression (OLCR) (A). The hallmark gene sets were obtained from the MSigDB database followed by GSEA based on the mRNA expression dataset from different GBM regions. The heatmap represents the normalized enrichment scores (NES) representing the degree to which a particular hallmark is significantly enriched (padj<0.05) in a specific GBM region. The color represents the value of NES where yellow and black indicate the highest (NES= 3.5) and lowest (NES= 0) NES values, respectively (B). Ki67 immunohistochemistry (IHC) of tumor core - superficial medial (C), anterior medial (D), tumor rim (E) and invasive margin (F) to estimate the fraction of proliferating cells in spatially distinct regions of GBM. The scale bar indicates 25 µm. CD31-IHC exploring tumour vascularity identifying the number of vascular structure in tumor core (G), tumor rim (H) and invasive margin (I). The scale bar indicates 100 µm. Heatmaps showing hierarchical clustering based on z-scored expression (Log_2_ TPM+1) of the leading edge genes of selected hallmarks - hypoxia (J), and inflammatory response/ TNFA signaling via NFKB (K). The leading edge genes representing the genes that contribute significantly to higher NES were identified from the GSEA output. Both the rows (genes) and columns (samples) were clustered using the Euclidian distance method. The column samples are further separated into different regions: Core (Tumor-core), Rim (Tumor-rim), Inv (Invasive margin), 5ALA sorted cells (5ALA+ and 5ALA-). The color code indicates the differential expression of leading-edge genes where red and blue represent higher and lower expressions, respectively. Some rows are labeled with the symbols of selected genes of interest.

To further investigate these enriched pathways, the leading edge genes contributing to the enrichment were identified for hypoxia, glycolysis, TNFA signaling via NFKB, and inflammatory response. Several genes associated with TNFA signaling via NFKB, and inflammatory response pathways showed an overlap; thus these two pathways were merged and common genes retained. A hierarchical clustering algorithm (Euclidian distance) was applied to the leading edge genes associated with hypoxia, glycolysis, and inflammatory response/ TNFA signaling via NFKB pathways (Supplementary Table S2), followed by the visualization of normalized (z-scored) gene expressions across different regions of 10 GBM patients as heatmaps (Figure 1J-K and Supplementary Figure S2G). Differential expression of these genes revealed that inter-tumor Core and Rim regions showed significantly higher relative expression of hypoxia-response related genes (Figure 1J). In contrast, both the unsorted Inv margin and the sorted invasive cells (5ALA+ and 5ALA-) showed significanlty lower relative expression of hypoxia-response genes in most of the patients (Figure 1J). A Kruskal-Wallis test confirmed the hypoxia-associated gene expression as significantly upregulated in tumor-core compared to the Inv margin, 5ALA+, and 5ALA- cells (Supplementary Figure S2I). Similarly, genes associated with glycolysis showed comparable differential regulation and were significantly upregulated in the Core and Rim regions relative to the Inv margin, 5ALA+, and 5ALA- cells (Supplementary Figure S2G and S2H). In contrast, TNFA signaling via NFKB, and inflammatory response genes exhibited significantly higher expression was in 5ALA+ and 5ALA- cells relative to all unsorted tumor regions (Figure 1K and Supplementary Figure S2J).

Next, we classified neural cell types to different GBM regions through enrichment analysis of gene signatures representing four neural cell types – Oligodendrocytes, Neurons, Astrocytes, and Cultured Astroglia (Supplementary Table S3). Core and Rim regions were enriched with all four cell types representing a heterogeneous cell population, whereas the Inv margin was mostly enriched with the neuronal cell type (Supplementary Figure S2K). Akin to Inv, 5ALA- cells were also enriched with the neuronal cell type (Supplementary Figure S2K). Interestingly, no enriched neural cell type signature was identified for 5ALA+ cells, indicating a likely evolution to a unique or hybrid cell-type signature that cannot be defined using canonical neural classifiers.

In summary, GSEA followed by hierarchical clustering revealed differential regulation of cancer-related and metabolic pathways in distinct GBM intra-tumor regions, suggesting adaptation to different microenvironmental selection pressures. 5ALA+ cells exhibited an upregulation of TNFA signaling via NFKB, and inflammatory response pathways, but could be classified by any known neural cell-type signatures.

### Disparate subtype enrichment within distinct GBM intra-tumor regions and 5ALA sorted cells

To further characterize spatially both distinct GBM regions and 5ALA sorted cells, we utilized enrichment analysis to identify GBM molecular subtypes (CL, NE, PN, and MES) previously described by Verhaak *et al.* ^10^ (Supplementary Table S4). A differential pattern of subtype enrichment was observed in spatially distinct GBM regions and 5ALA sorted cells; for instance, Core and Rim regions were mostly enriched with CL and PN subtypes (Figure 2A and Supplementary Figure S3A and B), whereas the NE subtype was highly enriched in the Inv margin (Figure 2A and Supplementary Figure S3C). 5ALA- cells were most closely associated with the NE and PN (Figure 2A and Supplementary Figure S3D). In contrast, 5ALA+ cells were highly enriched with the MES subtype (Figure 2A) with a high NES (2.10) and low adjusted p-value (2.3×10^-6^)

**Figure 2:**
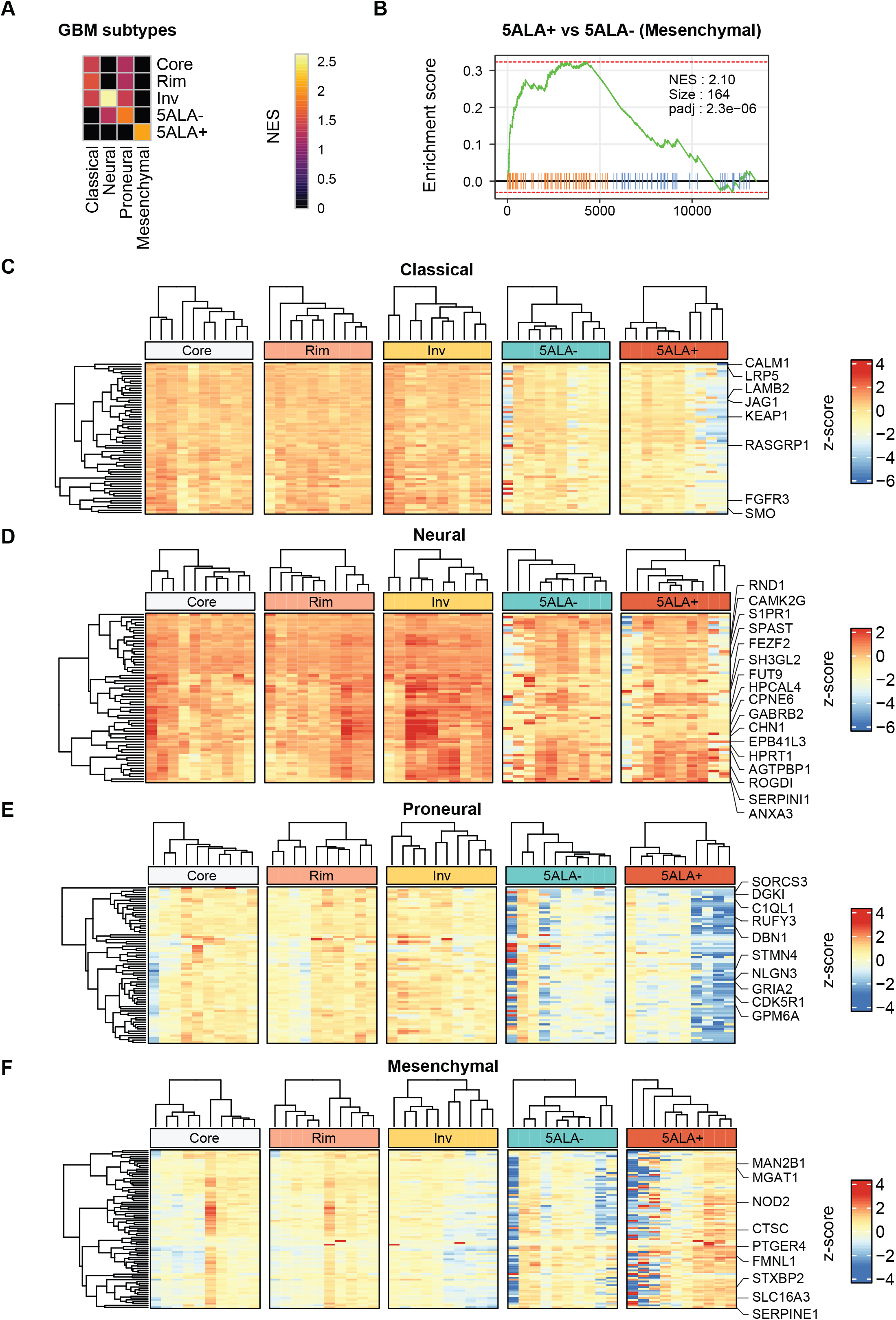
Enrichment of GBM subtypes in distinct GBM regions. Retrieval of gene-sets specifying different GBM subtypes was followed by GSEA. The NES of the significantly enriched (padj<0.05) GBM subtypes are shown in distinct GBM regions (Core, Rim, Inv) and 5ALA sorted cells (5ALA- and 5ALA+) (A). The color code indicates the differential NES values where yellow and black represent higher and lower normalized enrichment scores (NES) respectively. GSEA plot shows that the GBM mesenchymal subtype-specific genes are significantly enriched (NES: 2.1, padj=2.3×10^-6^) in 5ALA+ cells (B). Differential z-scored normalized expression (Log_2_ TPM+1) of the leading edge genes of GBM subtypes – Classical (C), Neural (D), Proneural (E), and Mesenchymal (F) - are shown as a heatmap. Column (samples) and row (genes) are clustered by the Euclidian distance method. The color code represents the differential expression of genes (Yellow: higher expression; Blue = lower expression).

(Figure 2B). The enrichment of the MES subtype is unique to 5ALA+ cells compared to all other regions and 5ALA- cells. To gain a deeper insight into the differential expression of subtype-specific genes, we identified the leading edge genes that contributed significantly to the enrichment of a subtype for a specific GBM region/cellular population (Supplementary Table S4). Hierarchical clustering analysis of the leading edge genes showed that CL-specific genes were highly expressed in the Core and Rim relative to 5ALA sorted cells (Figure 2C and Supplementary Figure S3E), whereas the Inv margin showed a higher expression of NE- and PN-specific genes across most of the patients, relative to all other regions and 5ALA sorted cells (Figure 2D, E and Supplementary Figure S3F and S3G). MES-specific genes exhibited a differential expression with relatively higher expression in the 5ALA+ cells (Figure 2F and Supplementary Figure S3H). Interestingly, a patient-specific differential expression was observed in 5ALA+ cells for MES-specific genes. For instance, 3/10 patients with variable expression patterns remained distinct whilst the remaining 7/10 patients were grouped into two separate clusters; the largest cluster comprising four patients showed the highest expression of MES-specific genes (Figure 2F).

Collectively, these results suggest that GBM molecular subtypes do not manifest uniformally throughout spatially distinct regions, but rather may vary in a region-specific manner. 5ALA+ cells were uniquely enriched with the MES subtype, with neither any intra-tumor region nor 5ALA- cells being enriched for MES.

### Inflammatory wound response, glycolytic/plurimetabolic, and mitochondrial gene signatures are enriched in 5ALA+ cells

We next applied gene-signatures characterizing two recently described cellular states - developmental and inflammatory wound response ^8^ and two metabolic states - mitochondrial (MTC) and glycolytic/plurimetabolic (GPM) ^9^ of GBM cells and performed GSEA in spatially-distinct GBM regions and 5ALA sorted cells (Supplementary Table S5). GSEA revealed a differential enrichment of these cellular and metabolic subtypes across GBM regions (Figure 3A). GPM and developmental subtypes were enriched in Core and Rim regions, with only the developmental subtype enriched in the Inv margin. Both GPM and MTC states were enriched in 5ALA+ and 5ALA- cells, indicating that similar metabolic states of 5ALA sorted cells may reflect the role of the shared infiltrative margin microenvironment. Among the metabolic states, GPM enrichment scores were higher in 5ALA+ cells whereas MTC enrichment score was higher in the 5ALA- cells (Figure 3A). Interestingly, inflammatory wound response was uniquely enriched in 5ALA+ cells (Figure 3A). For 5ALA+ cells, the highest enrichment score was observed for GPM (Figure 3B), followed by MTC (Figure 3C) and inflammatory wound response (Figure 3D). The leading edge genes of these distinct cellular and metabolic states were identified from GSEA output (Supplementary Table S5) and subjected to clustering analysis. A higher expression of GPM-associated genes in the 5ALA+ cells was observed for most of the patients (Figure 3E and Supplementary Figure S4A), whereas MTC genes were mostly upregulated in the 5ALA- cells (Figure 3F and Supplementary Figure S4B). Most of the genes associated with inflammatory wound response were highly upregulated in the 5ALA+ cells relative to other regions (5ALA+ *vs.* Core: p-value= 0.005 and 5ALA+ *vs.* Inv: p-value= 0.0006) and 5ALA- cells (Figure 3D, G and Supplementary Figure S4C).

**Figure 3:**
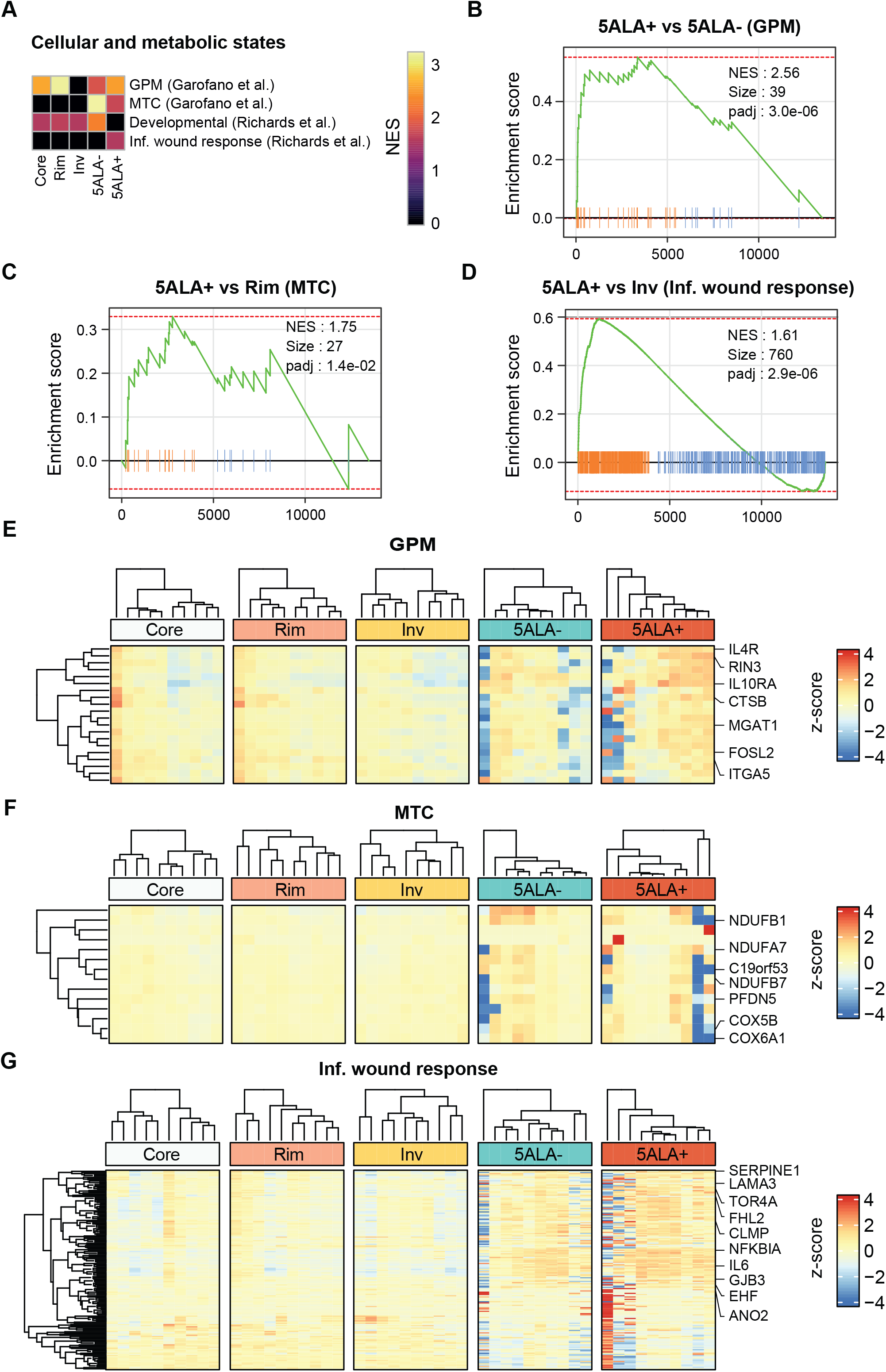
Enrichment of cellular and metabolic subtypes in distinct GBM regions. The gene-signatures of cellular states (Developmental and Inflammatory wound response) and metabolic states (Glycolytic - GPM and Mitochondrial - MTC) were retrieved and subjected to GSEA. Heatmap illustrating the normalized enrichment scores (NES) representing enriched cellular and metabolic states with (padj<0.05) in distinct GBM intra-tumor regions and 5ALA sorted cells (A). GSEA plots representing the enrichment of GPM-associated genes (B), MTC (C), and Inflammatory wound response (D) enriched in 5ALA+ cells compared to 5ALA- cells, Rim and Inv margin respectively. Heatmaps showing the differential expression of leading-edge genes of GPM (E), MTC (F), and Inflammatory wound response (G) in Core, Rim, Inv, 5ALA- and 5ALA+ cells. [Inf. wound response = Inflammatory wound response].

In summary, these results highlight the existence of differential cellular and metabolic subtypes throughout spatially distinct GBM regions and 5ALA sorted cells. Similar metabolic states in the 5ALA+ and 5ALA- non neoplastic cells were identified, whereas the enrichment of the inflammatory wound response pathway was unique to the 5ALA+ infiltrative tumor subpopulation.

### Identification of transcriptional networks controlling different cellular and metabolic states in 5ALA+ cells

In order to gain deeper insight into the transcriptional control regulating the cellular and metabolic gene-signatures enriched in 5ALA+ cells, we constructed a transcriptional network governing the leading edge genes of inflammatory response, TNFA signaling, MES, MTC, GPM, and inflammatory would response pathways. First, we identified TFs in the leading edge gene sets by using ENCODE TF annotation and in total, eight TFs were identified (KLF4, IRF8, NFKB1, NFKBIA, EGR2, REL, FOS, and FOSL2). These TFs and all leading edge genes were used as hubs and target genes, respectively in the ARACNE algorithm to identify the TF-target gene association significance. Only the TF- target gene associations with a p-value < 0.05 were considered. Based on the ARACNE output, a TF-target gene network was constructed and visualized by the Cytoscape tool (Figure 4A and Supplementary Table S6). All TFs (with the exception of FOS) showed a positive log_2_ fold change (FC) in 5ALA+ compared to 5ALA- cells, and among the positively regulated TFs, REL, IRF8, and NFKB1 exhibited the highest FCs. Most of the target genes (nodes) were also upregulated in 5ALA+ cells (Figure 4A). Among the TFs, FOSL2 showed the highest number (N = 103) of interaction (edges) with nodes, closely followed by IRF8 (N = 100) and NFKB1 (N = 85). The lowest number of target gene interactions was found for FOS (N = 35) whilst all other TFs exhibited higher (>70) target gene interactions. For TNFA signaling genes, the most prominent TF identified was NFKBIA, controlling 22 genes closely followed by EGR2 and KLF4, each controlling 21 genes (Figure 4B). The TFs controlling the highest number of inflammatory wound response genes were KLF4 (N = 33), IRF8 (N = 33), and NFKB1 (N = 32). For the Inflammatory pathway, the TFs controlling the most number of target genes, were IRF8 (N = 20), FOSL2 (N = 15), and REL (N = 14). The highest number of MES signature genes were likely to be controlled by FOSL2 (N = 30), IRF8 (N= 29) and NFKB1 (N=26), with FOLS2 also appearing to control seven MTC- and six GPM-associated genes. Eight GPM genes were identified to be controlled by NFKB1 and EGR2 (Figure 4B).

**Figure 4:**
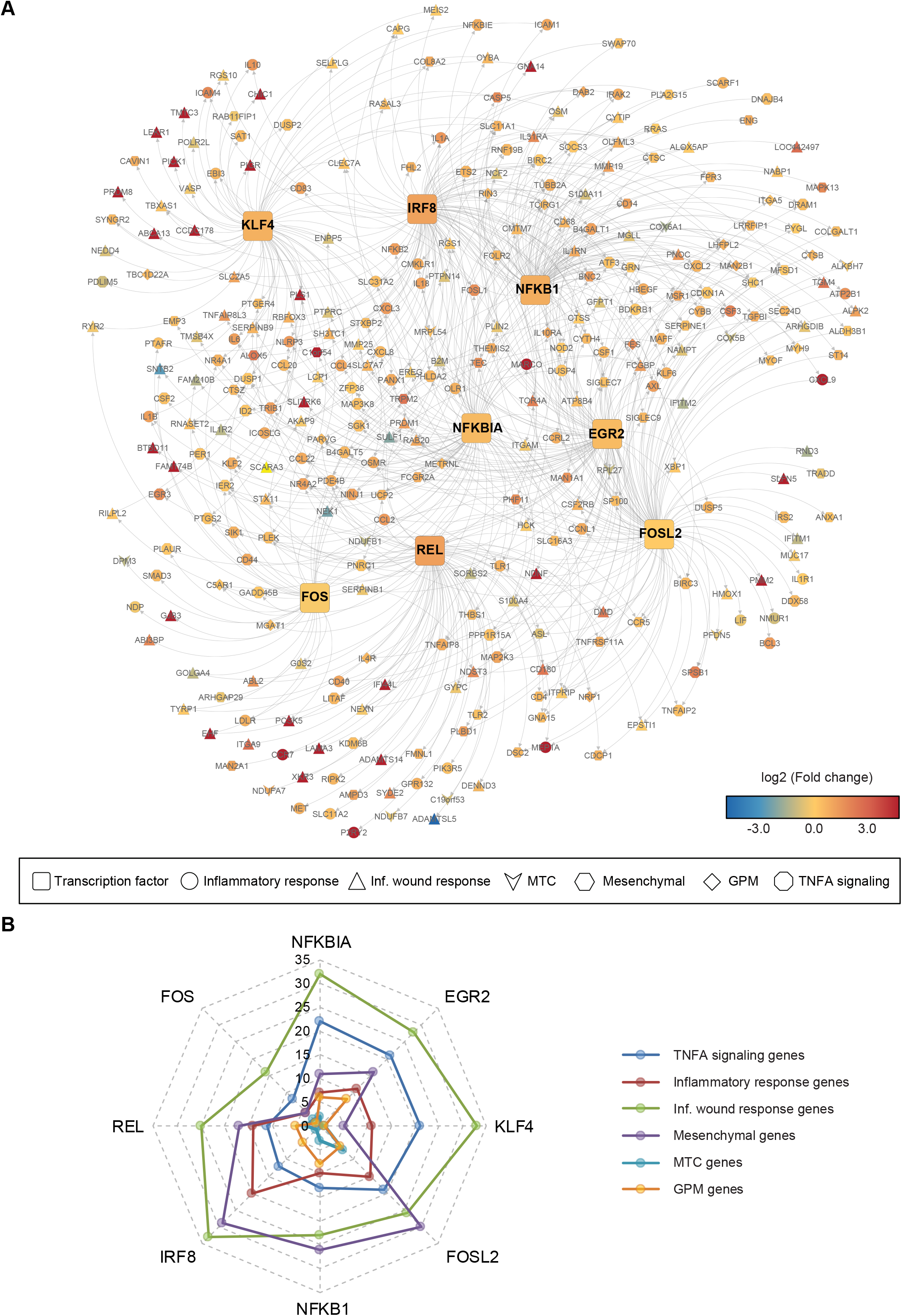
Construction of transcription networks upregulated in 5ALA+ cells. Transcription factors (TF) were identified in the leading edge genes that were enriched in 5ALA+ cells by using ENCODE TF-annotation. The association between TF and corresponding target genes (TG) was retrieved from the ENCODE database. A transcription network is constructed representing TF-TG association by using the ARACNe algorithm. The diagram illustrates that TGs (Nodes) associated with enriched gene sets (Inflammatory response, Inflammatory wound response, MTC, Mesenchymal subtype, GPM and TNFA signaling), are mutually controlled by different TFs (Hubs). The association between a node (TG) and a hub (TF) is shown as edges (grey curved lines) and nodes/hubs are color-coded according to the Log_2_ fold change (FC) values between 5ALA+ and 5ALA- cells (A), where red and blue indicate higher and lower FCs respectively. A radar plot representing TG (of different enriched subtypes) counts of the eight TF (B). The colored dots represent the TG count for a specific TF.

Overall, these results uncovered the transcriptional network controlling signature genes associated with enriched cellular and metabolic states in 5ALA+ infiltrative GBM cells. TNFA signaling and inflammatory wound response pathways are primarily controlled by the NFKBIA, EGR2, and KLF4 TFs, whereas the MES subtype was transcriptionally regulated by IRF8, NFKB1, and FOSL2. MTC and GPM subtypes were likely to be under transcriptional control of FOSL2 and NFKB1/EGR2, respectively.

### Higher exon and intron counts of the 5ALA+ enriched subtypes uncover transcriptional and post-transcriptional regulation

Exon-intron split analysis (EISA) was performed to determine the changes in pre-mRNA (intron) and mature-mRNA (exon) counts across distinct GBM regions and 5ALA sorted cells. We performed differential expression analysis to identify genes with altered exon and intron counts in 5ALA+ cells relative to tumor Core (Figure 5A), Rim (Figure 5B), and Inv margin (Figure 5C). A higher number of genes with significant intronic changes (Δintron) was observed in 5ALA+ cells compared to genes with significant changes in exon counts (Δexon) (Figure 5A-C). The higher number of genes with altered intron counts indicates a relatively strong transcriptional control regulating the genomic landscape of 5ALA+ cells. In contrast to other regions, a lower number of genes with significant Δexon (N = 140) and Δintron (N = 93) counts were identified between 5ALA+ *vs.* 5ALA- cells (Figure 5D). This lower number of genes with altered exon/intron counts between 5ALA sorted cells underscors a similar transcriptomic landscape of these cells, which may be induced by a shared infiltrative margin microenvironment. However, the contribution of technical factors inducing the experimental processes involving the isolation and preparation of 5ALA+/- cells for FACS analysis cannot be excluded. To functionally characterize the DEGs with altered intron and exon counts, hallmark enrichment analysis was performed. Oxidative phosphorylation genes with higher exon counts were highly enriched in 5ALA+ cells relative to Core, Rim, and Inv margin (Supplementary Figure 5A). This result was in agreement with our previous results, showing enrichment of the MTC subtype in 5ALA+ cells (Figure 3A). Genes with higher intron counts in contrast, were mostly associated with UV response apical surface and KRAS signaling pathways in 5ALA+ cells (Supplementary Figure 5A).

**Figure 5:**
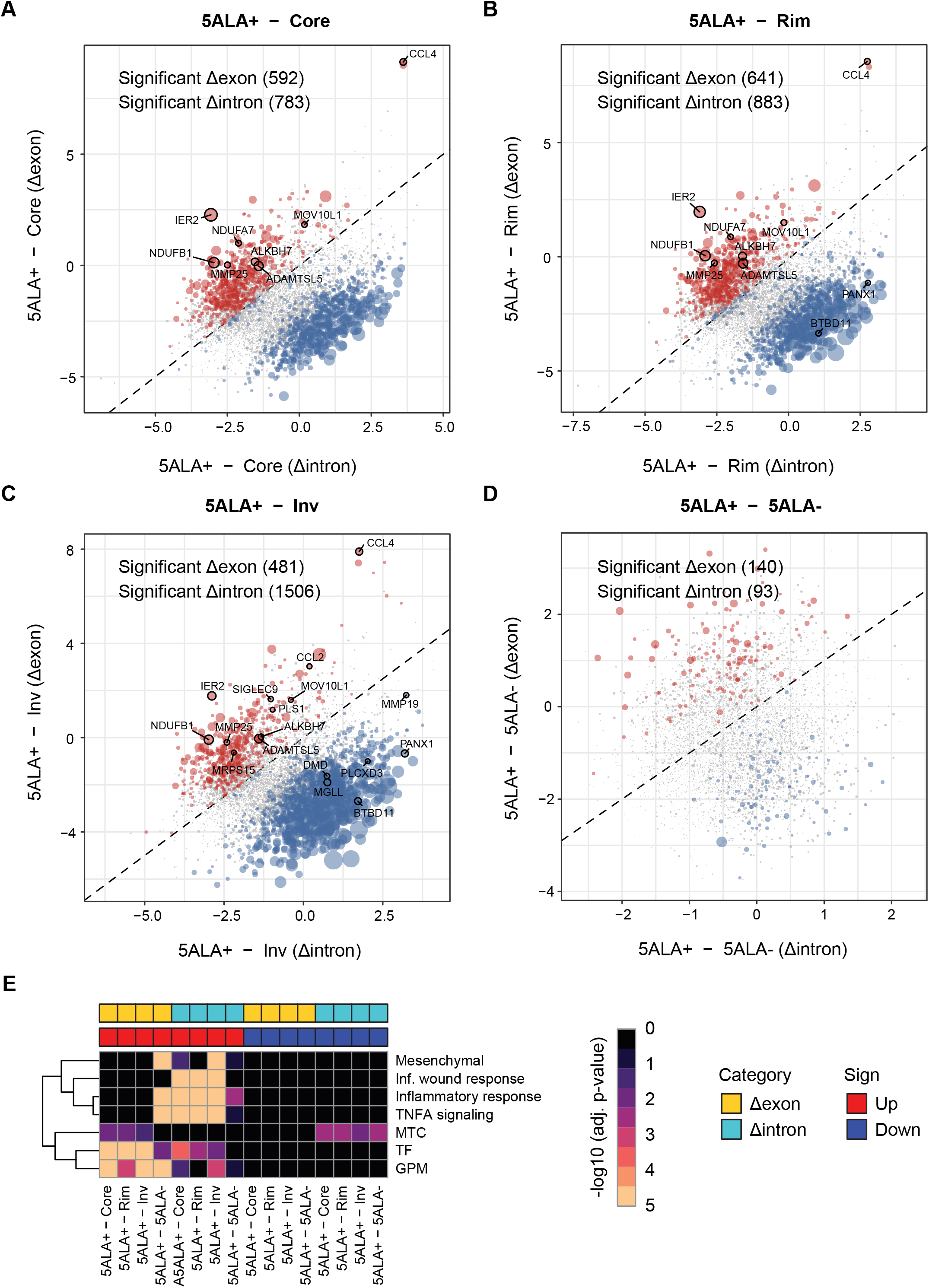
Exon-intron split analysis (EISA) of enriched cellular and metabolic subtypes in 5ALA+ cells. Scatter plot showing the gene-specific changes in exon (delta-exon) and intron (delta- intron) counts between 5ALA+ cells and tumor Core (A), Rim (B), Inv margin (C) and 5ALA- cells (D). X- and Y-axis represent the delta-intron and delta-exon, respectively. Color code represents the genes that showed significant regulation with adjusted p-value<0.05 (Blue: significant change in delta intron and Red: significant change in delta exon) between 5ALA+ cells and intra-tumor regions or 5ALA- cells. Heatmap showing the –log10 adjusted p-value of the hallmark enriched pathways (E). The samples representing the comparisons of 5ALA+ with other regions are given in columns (5ALA+ *vs.* tumor Core; 5ALA+ *vs.* tumor Riml 5ALA+ *vs.* Inv margin; and 5ALA+ *vs.* 5ALA-). Each row represents the different cellular and metabolic subtypes. Columns are divided into upregulated (Red) or downregulated (Blue) segments based on the exon and intrion regulation of genes in 5ALA+ cells. Finally, delta-exon (yellow) and delta-intron (Light blue) are annotated for each column. The color code in the heatmap is representative of –log10 p-values (Orange is highly significant and Black is not significant).

Furthermore, to investigate the transcriptional and post-transcriptional control of 5ALA+ cell specific cellular/metabolic gene-signatures and TFs, we obtained Δexon and Δintron counts of the leading edge genes representing enriched gene-signatures and upregulated TFs between 5ALA+ cells relative to unsorted regions and 5ALA- cells. Upon ranking the genes of each signature based on the Δexon and Δintron values, we performed pre-ranked GESA to investigate the enrichment of the gene-signatures (Supplementary Table S7). The results showed significant enrichment of MES subtype, inflammatory response and TNFA signaling pathways, GPM subtype, and TFs genes with upregulated Δexon counts in 5ALA+ cells (Figure 5E) relative to 5ALA- cells. In contrast, only inflammatory response genes with upregulated Δintron counts were enriched in 5ALA+ cells compared to 5ALA- cells (Figure 5E). Interestingly, a comparison of 5ALA+ cells with unsorted regions (Core, Rim, and Inv) mostly resulted in the enrichment of wound response, inflammatory response, TNFA signaling, and MES subtype with increased Δintron counts (Figure 5E). These pathways are likely to be transcriptionally restricted in 5ALA+ cells. Two enriched pathways (TFs and GPM) with upregulated Δintron counts were enriched in 5ALA+ cells relative to both unsorted tumor regions and sorted 5ALA- cells (Figure 5E) signifying transcriptional regulation controlling these genes.

Next, we identified significant DEGs with higher exon and intron counts in 5ALA+ cells. The highest number of DEGs were identified for wound response (N = 11) (Supplementary Figure S5B) followed by MTC (N = 5) (Supplementary Figure S5C) and TNFA signaling (N = 4) (Supplementary Figure S5D). Only one DEG was identified for the MES subtype (Supplementary Figure S5E) and inflammatory response pathway (Supplementary Figure S5F). For wound response, six genes (*IER2, PLSJ, MOV10L1, CCL2, MMP25, and ADAMTSL5*) exhibited increased Δexon and five genes (*MMP19, DMD, PLCXD3, MGLL, BTBD11*) showed increased Δintron in 5ALA+ cells (Supplementary Figure S6). All DEGs associated with TNFA signaling and MTC (except *PANX1*) exhibited higher Δexon counts in 5ALA+ cells (Supplementary Figure S6). *CCL2* associated with multiple subtypes – (wound response, inflammatory response, and TNFA signaling) with significantly higher (p-value<0.001) Δexon counts in 5ALA+ cells (Supplementary Figure S5D and S5F and S6L).

Collectively, these results decipher the transcriptional and post-transcriptional regulation of enriched cellular and metabolic gene-signatures in 5ALA+ infiltrative GBM.

### Higher stemness profile and enriched gene-signatures of 5ALA+ cells may contribute to the recurrence of GBM

Estimation of a patient-specific stemness index based on mRNA expression (mRNAsi) across spatially distinct GBM regions and 5ALA sorted cells was performed by the method described by Malta *et al.* ^34^. Hierarchical clustering (correlation-based) showed distinct mRNAsi profiles for 5ALA sorted (5ALA+ and 5ALA-) cell populations and unsorted Inv margin regions across 10 patients. Tumor Core and Rim regions exhibited similar but low mRNAsi (Figure 6A). Interestingly, differential stemness profiles were observed in a patient-specific manner. For instance, 5ALA+ cell populations of three patients (sample 34, 31, and 30) exhibited a higher stemness index compared to the remaining seven patients (Figure 6A). Similar differential stemness profiles across patients were observed for 5ALA- cells and other unsorted regions. With the highest stemness profile, 5ALA+ cells showed a distinct stemness profile relative to all unsorted tumor regions (Figure 6A and B). Although the stemness profile of 5ALA+ cells was higher than 5ALA- cells and unsorted Invasive margin tissue, the differences were not statistically significant (Figure 6B). Interestingly, TCGA-GBM samples (representing the GBM core) showed a relatively lower mRNAsi index compared to 5ALA+ cells. Stratification of the TCGA-GBM samples into molecular subtypes revealed a lower stemness index of the GBM subtypes compared to both 5ALA+ and 5ALA- cells (Figure 6B). To identify the association between gene-signatures and the mRNAsi of the 5ALA+ cells, we retrieved the stemness-associated genes (*KLF4, MYC, CTNNB1, EPAS1, EZH2, KDM5B, NES, TWIST1, ABCG2, CD34, CD44, NANOG, PROM1, ZFP42, and ZSCAN4*) as reported by Malta *et al.* ^34^ and analyzed the correlation between mRNAsi and expression of these genes across the 10 patients (Supplementary Figure S7A). Only two genes - *CD44* and *ZSCAN4* - showed a positive correlation between mRNA expression and mRNAsi (Supplementary Figure S7A). Next, we obtained 13 known brain cancer and 13 stem cell markers from the cell marker database (http://biocc.hrbmu.edu.cn/CellMarker/help.jsp) ^47^ and investigated the correlation between mRNAsi and mRNA expression. In this case, two genes - *ITGA6*, *SLITRK6* and *SOX9* - showed a positive correlation (Supplementary Figure S7B). However, when the expression levels of these genes were analyzed, *CD44, ZSCAN4, SLITRK6,* and *SOX9* showed low expression in 5ALA+ cells (Supplementary Figure S7C and S7D).

**Figure 6:**
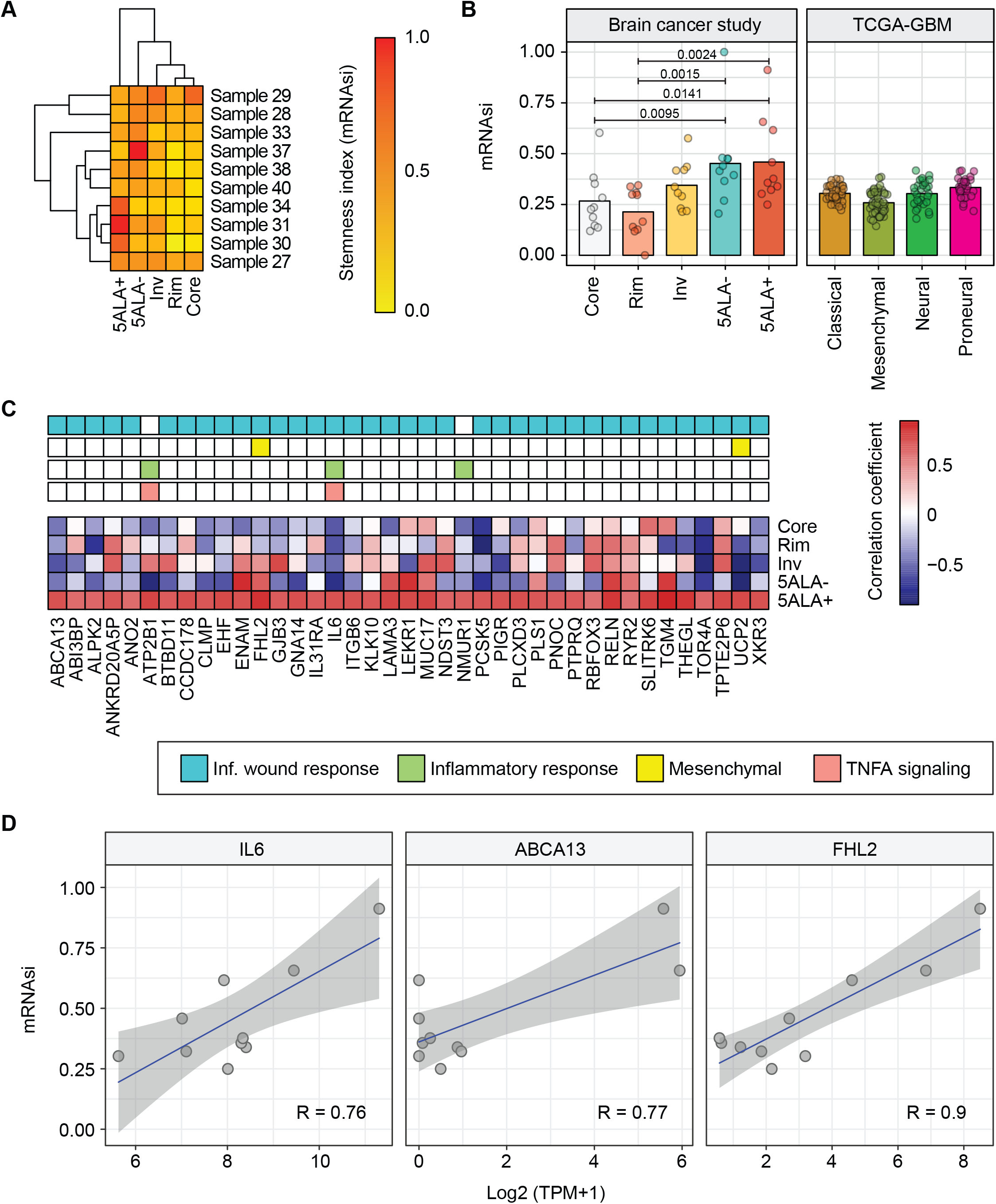
Higher stemness index of 5ALA+ cells defines ‘*GBM Infiltrative Stem’*. mRNA-based stemness index (mRNAsi) values across distinct GBM regions (Core, Rim, Inv, 5ALA- and 5ALA+) for each patient are represented in a heatmap. The values are scaled between 0 (low) and 1 (high) (A) and are indicated by the gradient color code. Row (samples) and columns (GBM regions) are clustered by using a correlation algorithm. Comparison of the mRNAsi values according to brain regions (left) and TCGA-GBM samples (right) are shown as bar diagrams (B). The TCGA-GBM samples were pre-stratified according to GBM subtypes as described by Verhaak et al. ^10^ Kruskal-wallis tests showed a significantly higher mRNAsi in 5ALA+ cells compared to Core and Rim, with p-values shown. Correlation (Pearson correlation coefficient) values between the mRNAsi and mRNA expression of selected genes are shown (C). The genes that showed a significant (p-value < 0.05) positive correlation with mRNAsi for 5ALA+ cells for each patient were selected. Correlation values are indicated by the color code where red and blue represent positive and negative correlation respectively. The scatter plots show the correlation of mRNAsi values and mRNA expression of three selected genes (*IL6*, *ABCA13*, and *FHL2*) (D).

To further characterize the stemness associated gene-signature, eight previously published stemness associated gene sets representing Consensus Stemness (Shaat *et al.*) ^31^, Human embryonic stem cell - HuESC (Bhattacharya *et al.*) ^30^, Stem cell (Palmer *et al.*) ^27^, Myc induced genes (Kim *et al.*) ^32^, Embryonic stem cell - ES1 (Ben-porath *et al.*), Sox2 induced genes (Ben-porath *et al.*) ^29^, NANOG induced genes (Ben-porath *et al.*) ^29^ and Epithelial Atypical squamous cells (ASC) (Richards *et al.*) ^8^ were retrieved. GSEA showed that most of the gene-sets were highly enriched in the Core, followed by Rim and Inv margin. 5ALA- cells were enriched in Consensus Stemness, HuESC, and Myc gene sets. Interestingly, none of these gene-sets were enriched in 5ALA+ cells (Supplementary Figure S7E and Supplementary Table S8). These results all together underscored the uniqueness of the 5ALA+ cells implying that previously published canonical stemness-associated gene sets may not be accurate, nor informative when characterizing the unique stemness properties of 5ALA+ infiltrative GBM.

Realizing the uniqueness of the 5ALA+ cells, we obtained the six gene-signatures that were enriched in 5ALA+ cells, in addition to the correlation analysis between stemness and mRNA expression. A total of 39 genes mostly associated with inflammatory wound response, MES subtype, inflammatory response, and TNFA signaling pathways, showed a significant positive correlation with mRNAsi, and constitutes a holistic dataset, which we term „*GBM Infiltrative Stem*‟ (Figure 6C and Supplementary Table S9). The highest correlation was observed for the gene - Transglutaminase 4 (*TGM4*; R= 0.95). The impact of *GBM Infiltrative Stem* on survival of GBM patients from the TCGA cohort were investigated by a cox regression model. Three genes - Interleukin 6 (*IL6*), ATP Binding Cassette Subfamily A Member 13 (*ABCA13*), and Four and a half LIM domains protein 2 (*FHL2*) - were associated with poor survival outcome and a high correlation with mRNAsi (FHL2, R= 0.90; IL6: R= 0.76; and ABCA13, R= 0.77) (Figure 6D).

Overall, these findings encourage us to define *GBM Infiltrative Stem* as a 5ALA+ gene signature positively correlated with inflammatory would response, MES subtype, inflammatory response and TNFA signaling gene-signatures.

### Enriched cellular and metabolic subtypes of 5ALA+ are associated with poor overall survival

Six 5ALA+ enriched cellular and metabolic gene-signatures (TNFA signaling, inflammatory response, MES subtype, inflammatory response, GPM and MTC) were subjected to univariate cox regression analysis by using TCGA-GBM patient survival data followed by the Hazard ratios (HRs) calculation for individual genes. Genes with an HR of > 1 and a p-value < 0.05 were considered to be associated with risk (Supplementary Table S10). Cox regression model revealed a significant risk associated with genes defining all subtypes except MTC. The highest number of risk-associated genes were identified for the MES subtype (N = 42), followed by inflammatory wound response (N = 30), TNFA signaling (N = 27), and inflammatory response (N = 23) (Supplementary Table S10). Chemokine *CCL2* with significantly higher exon count in 5ALA+ cells (Supplementary Figure 5D) exhibited a significant impact on the survival outcome of TCGA-GBM patients (HR: 1.1, min: 1 – max: 1.3, p-value= 0.01) (Supplementary Table 10). Three genes (*IL6*, *ABCA13*, and *FHL2* that exhibited high positive correlation with mRNAsi were also associated with poor survival outcome.

To evaluate the combined impact of risk-associated genes on survival outcome, TCGA-GBM patients were classified into high- and low-expression cohorts by using a median expression threshold for each subtype to determine overall survival (OS) (Figure 7 A-E) and disease-free survival (DFS) (Supplementary Figure S8A-G). Kaplan–Meier curves showed that all 5ALA+ enriched gene-signatures were associated with poor DFS, with the exception of the MTC subtype (Supplementary Figure S8A-G). MES (Figure 7C), inflammatory wound response (Figure 7D) and GPM (Figure 7E) were associated with poor OS where the highest significance was observed for the MES subtype (p-value = 0.0084). The highest significance (lowest LogRank p-value) for poor DFS outcome was associated with a high-expression group of inflammatory response (Supplementary Figure 8B), followed by inflammatroy wound response (Supplementary Figure 8D) and MES (Supplementary Figure 8C) gene-signatures.

**Figure 7:**
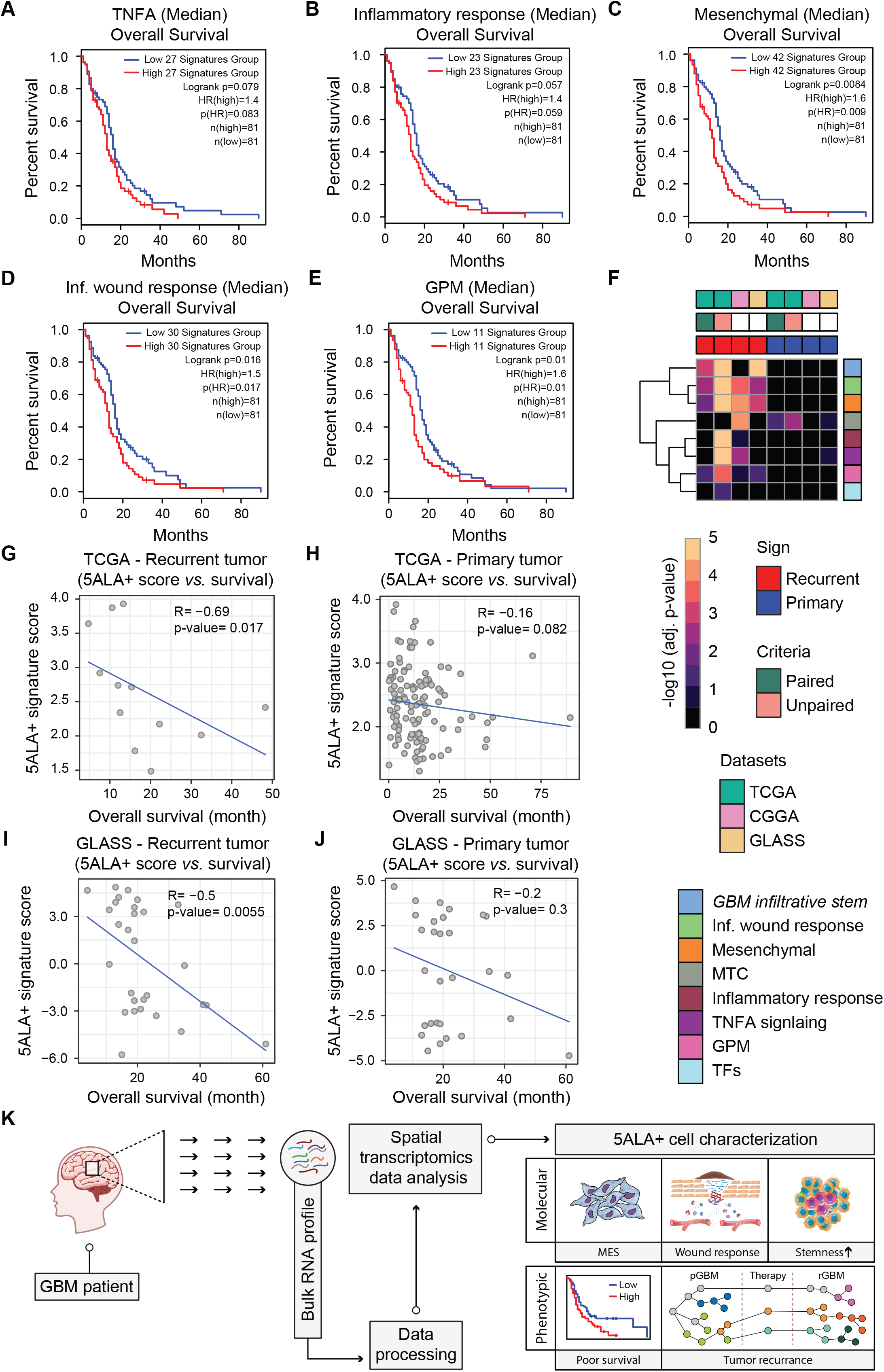
Association of 5ALA+ gene-signatures with overall survival and recurrence in GBM patients. Based on the median combined mRNA expression of five different cellular and metabolic subtypes (TNFA signaling, Inflammatory response, Mesenchymal subtype, Inflammatary wound response and GPM), TCGA-GBM patients were stratified into high- (N = 81) and low-expression (N = 81) groups. Only the genes that showed a significant association with GBM (hazard ratio>1 and p-value<0.05) in univariate analysis were considered for each subtype. The number of genes in each subtype is shown: TNFA signaling (N = 27), Inflammatory response (N = 23), Mesenchymal subtype (N = 42), Inflammatory wound response (N = 30) and GPM (N = 11). Log-rank p-value indicates the significant difference in survival outcomes between high- and low-expression cohorts. Kaplan-Meier curves represent the overall survival outcomes of TNFA signaling (A), Inflammatory response (B), Mesenchymal (C) Inflammatory wound response (D) and GPM (E) gene signatures respectively. Heatmap showing the –log10 adjusted p- value of the enriched gene signatures across recurrent and primary GBM tumors (F). The conditions representing the comparisons between recurrent and primary GBM tumors from TCGA and CCGA are given in columns (TCGA - Recurrent *vs.* Primary and CCGA - Recurrent *vs.* Primary). GBM data from TCGA has been analyzed in a paired and unpaired manner. Columns are divided into upregulated (Red) or downregulated (Blue) segments based on the regulation of genes between Recurrent *vs.* Primary samples. Each row represents the different gene signatures. The color code in the heatmap is representative of –log10 p-values (Orange is high and Black is low). Scatter plots representing the correlation between 5ALA+ gene-signature scores (combined normalized enrichment score for Mesenchymal and Inflammatory wound response for each patient) and overall survival (months) in recurrent and primary patients from TCGA (G and H) and GLASS (I and J) cohorts. Spearman correlation coefficient (R) and p- values are shown.

To explore the impact of the 5ALA+ enriched gene signatures on the recurrence of GBM, GSEA was performed on RNA-seq data obtained from primary and recurrent IDH wild-type patients of TCGA, CGGA and GLASS cohorts (Supplementary Table S11). First, a hallmark enrichment anlaysis between recurrent and primary samples revealed that most of the enriched pathways were detected in recurrent tumors. Primary tumors only exhibited marginal enrichment of pathways such as Hypoxia, DNA repair, E2F targets, whereas a diverse array of pathways including Oxidative phosphortlation, MYC targets, fatty acid metabolism, IL6/JAK/STAT signaling, Epithelial-Mesenchymal transition, TNFA signaling via NFkB and inflammatory response, were enriched at recurrence and where the two latter pathways were also enriched in 5ALA+ cells (Supplementary Figure S9). TCGA harbours an unequal sample distribution between primary (N = 154) and recurrent (N = 13) tumor samples. Among the thirteen recurrent GBM samples, six were paired with primary tumor data. Therefore paired and unpaired analyses were performed for TCGA (Supplementary Table S11). For CGGA cohorts primary and recurrent tumor samples are unpaired, however for GLASS cohorts the primary and recurrent samples are paired. Enrichment analysis revealed an intriguing phenomenon where 5ALA+ gene-signature enrichment was observed in recurrent but not in primary tumors. For instance, MES and inflammatory wound response were enriched in recurrent tumors across TCGA, CCGA and GLASS cohorts (Figure 7F). *GBM Infiltrative Stem* was enriched in recurrent GBM of TCGA and GLASS cohorts but not in CGGA (Figure 7F). Compared to recurrent tumors, only MTC gene signatures were marginally enriched in the primary tumors of TCGA (Figure 7F). To further investigate the impact of unique 5ALA+ gene-signatures on the survival of recurrent and primary GBM patients, ssGSEA was performed in recurrent and primary tumors separately for two unique gene-signatures – MES and inflamamtory wound response. By combining the normalized enrichment values, a combined 5ALA+ gene-signature socre was calculated for each tumor sample, followed by correlation analysis with survival data of recurrent and primary patients. For recurrent tumors of TCGA (Figure 7G) and GLASS (Figure 7I), 5ALA+ gene-signature scores exhibited a significant negative correlation with survival, whereas primary tumors did not show a significant correlation with survival (Figure 7H and J).

In conclusion, the enrichment gene signatures of 5ALA+ cells were associated with poor survival and recurrence in GBM, signifying the probable impact of these signatures in tumor progression and interval to disease recurrence (Figure 7I).

## Discussion

We have revealed distinct cellular, stemness and metabolic gene-signatures in spatially delineated intra-tumor GBM regions and 5ALA fluorescence-sorted invasive margin cells. Enriched gene-signatures in 5ALA+ cells, including our newly defined *GBM Infiltrative Stem*, are uniquely associated with poor survival and tumor recurrence, implicating 5ALA+ cells as accurate proximates of GBM residual disease post-surgery.

Previously, it has been hypothesized that 5ALA fluorescence beyond the T1 enhancing region on magnetic resonance imaging (MRI), represents a unique microenvironment contributing to molecular signatures distinct to tumor Core and Rim ^2^. Corroborating this hypothesis, we showed that pro-proliferative pathways (e.g. mitotic-spindle, G2M checkpoint, mTOCR1 signaling and E2F targets) were highly enriched in the Core but absent in the infiltrative margin (Figure 1B). Moreover, upregulation of glycolytic and angiogenesis pathways coupled with an absence of oxidative phosphorylation in the Core and Rim (Figure 1B) signifies a hypoxia-induced Warburg effect. In contrast, the Hypoxia-response pathway was absent and oxidative phosphorylation pathway upregulated, in both the unsorted invasive margin and 5ALA+ and 5ALA- sorted cells, implying a lack of hypoxic pressure in the microenvironment harbouring residual, infiltrative disease (Figure 1B). This encourages reconsideration of delivery carriers which are designed to trigger the release of therapeutic payloads only in low pH/reducing conditions.

5ALA+ cells showed a unique immune-system associated molecular signature including TNFA signaling via NFKB and inflammatory response (Figure 1B and 3A) relative to 5ALA- cells. This corroborates with our previously reported upregulation of inflammatory response and downregulation of hypoxia response genes in 5ALA+ cells ^14^. This refutes the notion that certain cellular states are artifactually established due to the process of FACS, and supports the claim that these states reflect the biology of infiltrative margin residual disease. Enrichment of the NEU cell type in both unsorted Inv and 5ALA- cells, indicates that normal neural cells constitute the majority of the infiltrative margin, consistent with previous findings^48^. One of the striking features of 5ALA+ cells was the lack of molecular identity to canonical neural cell types, underscoring the unique transcriptional landscape infiltrative GBM. Using a microarray-based study, Bonnin *et al.* also showed that 5ALA+ tumor tissue failed to exhibit molecular signatures of any known cell types ^49^.

Although GBM was originally classified into four molecular subtypes – PN, NE, CL and MES - ^10^, emerging evidence supports molecular subtype plasticity and subtype switching to MES in particular, being associated with recurrence and poor survival outcome ^11^. Recently, Minata *et al.* hypothesized that the switch of PN to MES subtype may assist the tumor cells by providing a survival advantage ^13^. To extend this hypothesis we show that the plasticity of GBM molecular subtypes is not restricted to recurrence, but can manifested in a region and cell-type-specific manner. Unsorted Core and Rim were highly enriched with CL and PN subtypes (Figure 2A), and unsorted Inv enriched with NEU (Figure 2A). 5ALA+ cells in contrast, were uniquely enriched with the MES subtype (Figure 2B). This finding supports a hypothesis that an infiltrative MES subtype may drive GBM recurrence. Interestingly, the link between a MES gene-signature and decreased tumor purity has been established as a common theme across different cancers ^50, 51^. One of the prominent features of MES is the association with immune-related pathways and the lower purity score in comparison to PM and CL, highlighting the possible infiltration of non-neoplastic and immune cells into MES ^11, 52^. In agreement with our previous results ^14^, the current study shed light on the association of higher MES gene expression and low tumor purity, by identifying a 5ALA+ MES gene-signature which constitutes only a minor fraction of the invasive margin.

Recently, by employing scRNA-seq and genome-wide CRISPR screening, Richards *et al.* extensively characterized the cellular phenotypes of GCSs and identified high inter- and intra-GSC transcriptional heterogeneity independent of DNA somatic alterations ^8^. Rather, the authors showed that GSCs harbor a transcriptional gradient ranging from neural development to wound response. However, they identified fewer wound response-like cells compared to neural developmental-like cells in primary tumor core regions and concluded that the transcriptional-gradient model can help global transcriptional heterogeneity of infrequent tumor-initiating cells in the GBM Core ^8^. In this study, we revealed an enriched neural developmental signature in spatially distinct GBM regions and 5ALA- cells, whereas a wound response signature was uniquely enriched in 5ALA+ cells. These findings corroborate evidence showing a lower number of tumor-initiating wound response-like cells in GBM ^53^. Another recent study identified MTC and GPM metabolic states traits by integrating multiple single cells and bulk tumor transcriptome datasets ^9^. The authors concluded that the MTC state that relies on oxidative phosphorylation for energy production was associated with the most favorable clinical outcome, whilst the GPM state mediated by aerobic glycolysis and amino acid/lipid metabolism, is linked to poor patient outcome ^9^. We provide evidence of GPM enrichment GBMCore and Rim but not Inv margin. Interestingly, both GPM and MTC states were enriched in 5ALA+ and 5ALA- cells, reflective of a shared microenvironment in controlling the metabolic states of both infiltrative tumor and non-neoplastic cells.

To better understand transcriptional regulatory mechanisms that underlie the upregulation of MES and wound response associated genes in 5ALA+ cells, we employed EISA – a method that can reliably distinguish between transcriptional and post-transcriptional control ^18^. Upregulation of exonic levels for MES and inflammatory wound-response genes in 5ALA+ relative to 5ALA- cells implied that these genes were under active post-transcriptional control. Previously, a mass-spectrometry-based study has shown that altered expression of nuclear regulatory proteins controlling transcription and post-transcriptional processes may drive GBM invasion ^54^. In addition to the differential abundance of nuclear proteins, the role of the microenvironment in the regulation of post-transcriptional processes including splicing and translational control cannot be ruled out ^55^. The common microenvironment shared by the 5ALA+ and 5ALA- cells argues against a direct role of the microenvironment to increase the exonic counts of the MES and inflammatory wound response genes only in 5ALA+ cells. However, the unique genetic background and cell lineage of the 5ALA+ cells may contributemicroenvironmental interaction distinct from that of normal neural cells residing in the infiltrative margin.

The transcriptional regulatory network that governs GBM transition to MES and recurrence has not been elucidated, partly due to an inability to identify and characterize rare GBM sub-populations exhibiting MES. The transition of GSCs to MES was reported to be dependent on TNFA signaling via NFkB pathway ^13^, which was upregulated in 5ALA+ cells in our study. Enrichment of MES both in the 5ALA+ subpopulation and recurrent tumors, offers a unique opportunity to explore 5ALA+ transcriptional networks further, to eludicate biomarkers predictive of recurrence interval and to identify putative molecular therapeutic targets to initiate treatment in advance of recurrence..

Interestingly, among the TFs, two NFkB family members - NFKB1 and REL - in addition to the inhibitor of NFkB-REL complex NFKBIA, were upregulated in 5ALA+ cells. These seemingly opposing factors may establish a delicate balance between inflammatory and anti-inflammatory pathways that are required to maintain a chronic and persistent low level of inflammation, further boosted by the infiltration of anti-inflammatory and regulatory immune cells ^56^. MES genes were mostly controlled by IRF8, NFKB1, and FOSL2. The upregulation of IRF8 in 5ALA+ cells was surprising, since it is considered a myeloid-lineage specific master TF exclusively restricted to the hematopoietic lineage ^57^. Moreover, expression of IRF8 was assumed to be repressed in neural stem cells ^58^ which raises the question of why and how 5ALA+ cells were able to express IRF8. In a recent study, Gangoso *et al.* shed light on this conundrum, by performing RNA-seq analysis of patient-derived GSC cultures, and based on differential immune-induced gene expression, identified two distinct subtypes – non-mesenchymal immune signature (Non-MESImm) and mesenchymal-immune signature (MESImm) ^58^. Remarkably, an upregulation of interferon regulatory factor family members in MESImm-enriched GSCs (including IRF8) was observed, suggesting that GSCs have the capacity to hijack myeloid-specific transcriptional modules to evade the immune response and chemokine expression such as CCL2 ^58^. As a mechanistic explanation of how MESImm GBM subtype cells are able to express IRF8 and CCL2, the authors observed hypomethylation at specific CpG islands associated with IRF8 and CCL2. These findings signify a possibility of epigenetic alteration reshaping the transcriptomics landscape of 5ALA+ cells to express IRF8 and CCL2. In agreement with the current results, FOSL2 was previously confirmed as a contributing factor of MES acquisition ^59^. Inflammatory wound response is evidently also controlled by IRF8, in addition to KLF4 and EGR2. KLF4 represents one of the key stemness factors in GSCs, and has been implicated in the survival of GSCs and GBM recurrence ^60^. Similarly, EGR2 was previously shown to promote glioma invasion ^61^.

The higher mRNA-based stemness index (mRNAsi) of 5ALA+ cells (which we have defined *GBM Infiltrative Stem*), relative to canonical stemness signatures of intra-tumor GBM regions and TCGA GBMs, urges pragmatic reflection on GSC biology to date. We reveal a higher mRNAsi in the GBM infiltrative margin relative to GBM Core and Rim, which may be explained by the presence of a higher number of neural progenitor cells in the GBM Inv margin. Previously, we reported a distinct molecular composition of the heterogeneous GBM infiltrative margin characterized by higher stem cell marker expression compared to GBM Core ^2^. The current study gained a deeper insight into the heterogeneity of the GBM Inv margin by extending the hypothesis that molecularly distinct 5ALA+ cells exhibit the highest stemness index. As non-neoplastic 5ALA- cells also exhibited a higher stemness profile, the microenvironment composing the GBM infiltrative margin is a probable determinant in the acquisition of stemness. Nevertheless, the impact of the microenvironment on transcriptional regulatory networks governing cancer stemness is not fully understood. Miranda *et al.* investigated the link between stemness and immune cell infiltration across 21 cancer types including GBM ^28^, demonstrating a negative correlation between cancer stemness and anti-tumor immune response and immunosuppressive pathways, independent of increased mutation load, cancer-antigen expression and intratumoral heterogeneity ^28^. Our observation of IRF8 and CCL2 upregulation in 5ALA+ cells may contribute to such immunosuppressive pathways by recruitment of regulatory immune cells.

Lastly, we associate 5ALA+ gene signatures with GBM recurrence. Gene-signatures representing stemness, MES and inflammatory wound response were enriched in recurrent GBM tumors relative to primary tumors and were associated with poor survivaloutcome. Recently, Varn *et al.* showed that MES transition of GBM is linked to the presence of a distinct myeloid cell state characterized by unique ligand-receptor interactions with malignant cells ^12^. This finding is consistent with our results showing MES subtype-enriched 5ALA+ cells may serve as the source of IRF8-mediated signaling, leading to the production and secretion of chemokines such as CCL2, which may in turn act as a chemo-attractive agent to recruit myeloid cells into the GBM infiltrative margin.

Overall, our results comprehensively revealed unique molecular characteristics of the 5ALA+ sub-population(s) within the GBM infiltrative margin and underscored the possibility that interaction of 5ALA+ cells with this microenvironment may be a critical determinant for survival outcome and GBM recurrence. Our findings encourage the neuro-oncology research community to the prioritise this infiltrative GBM subpopulation(s) for both mechanistic pre-clinincal modelling and to predicate and expedite next-generation molecular targeted drug screening.

---

**Supplementary Figure S1.**
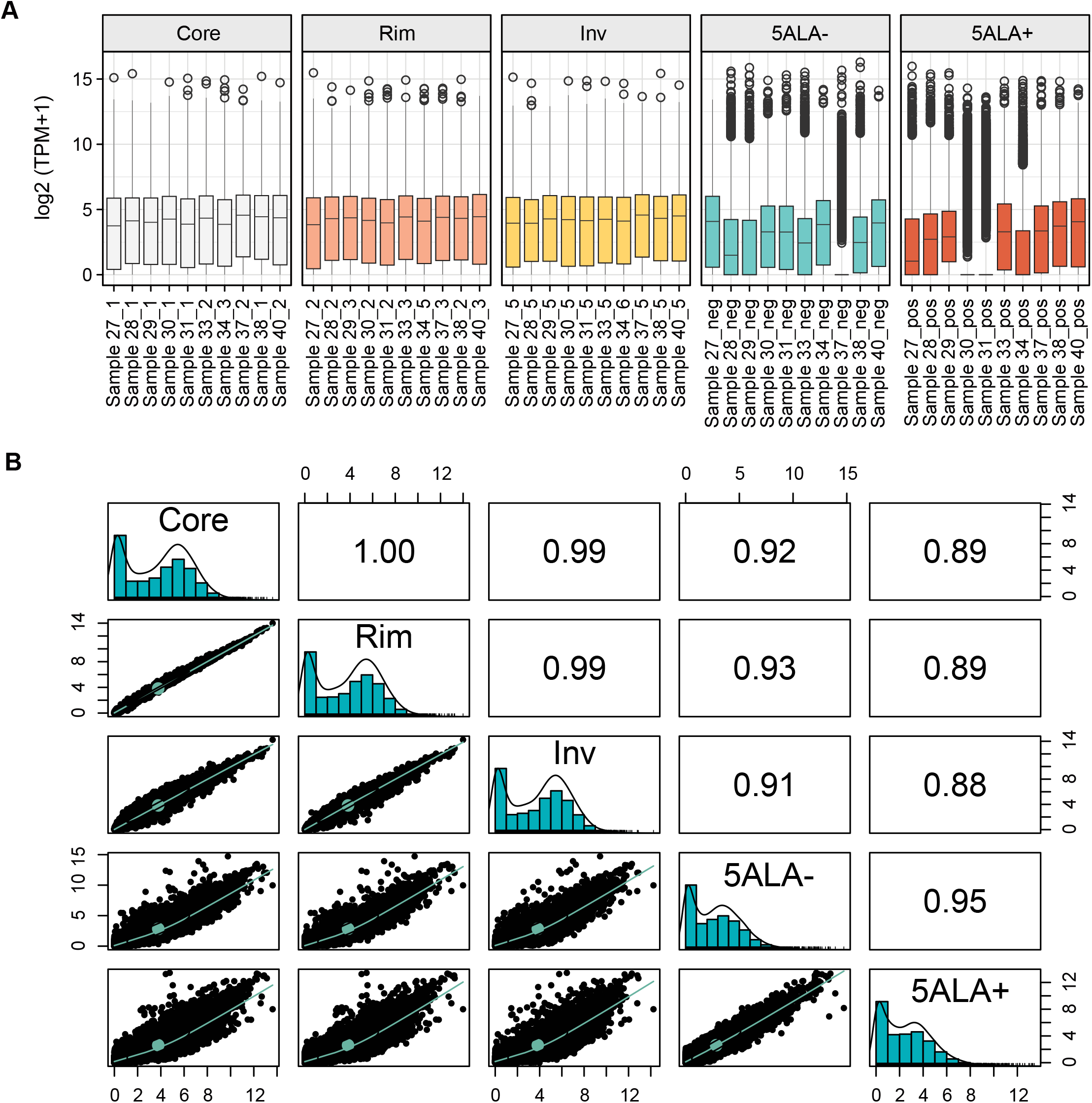
The boxplot of log normalized TPM expression values of 50 samples from 10 patients divided into five brain regions (Core, Rim, Inv, 5ALA-, 5ALA+) (A). The scatter plot matrix shows the histogram and correlation coefficients of all related brain regions. The lower triangle boxes represent the pairwise scatter plots, diagonal boxes show the brain regions with the distribution of the values for those regions, and the correlation coefficients are presented in the upper triangle boxes (B).

**Supplementary Figure S2.**
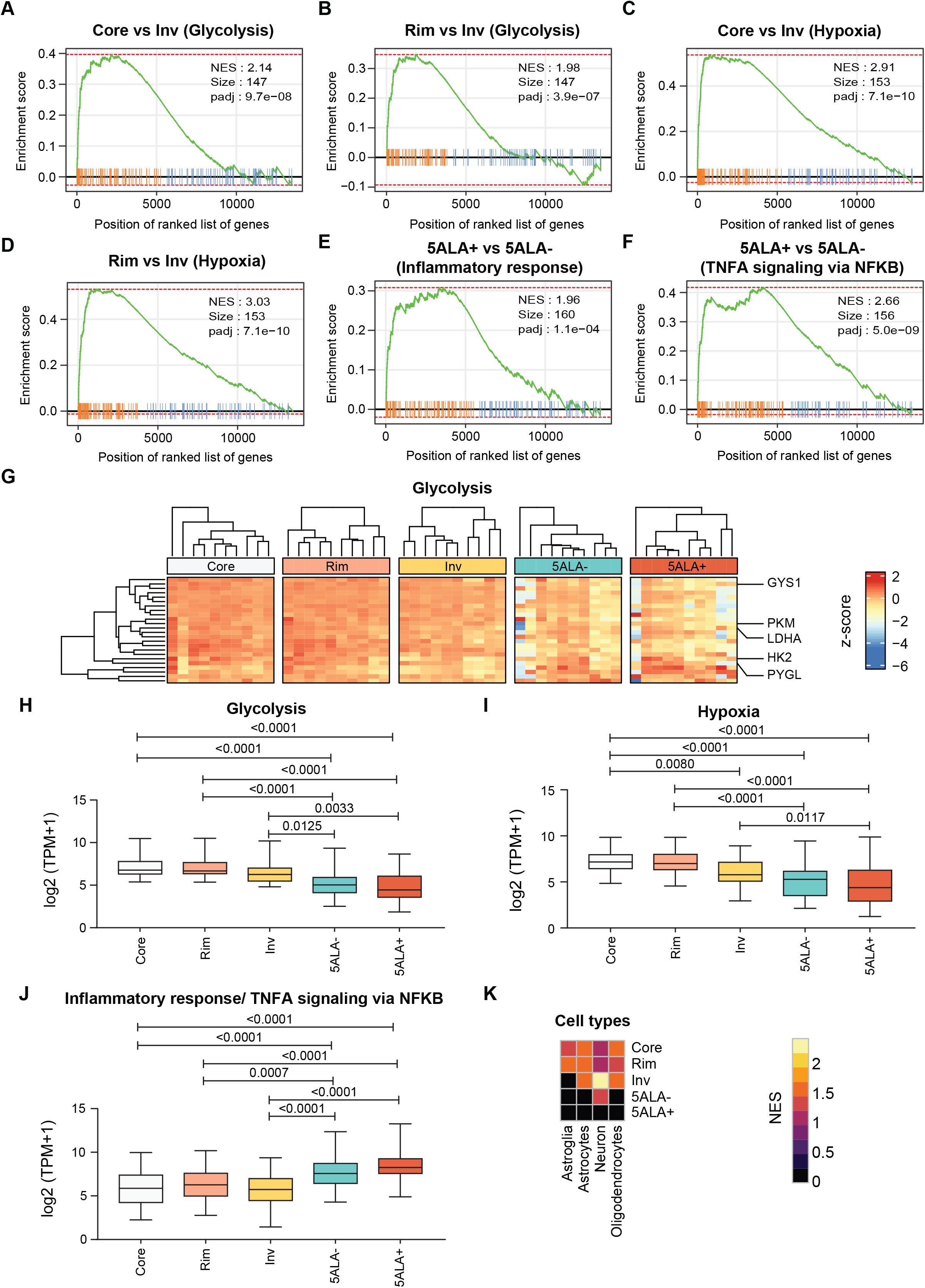
GSEA enrichment plots of representative gene sets from Glycolysis (Core *vs.* Inv) (A), Glycolysis (Rim *vs.* Inv) (B), Hypoxia (Core *vs.* Inv) (C), Hypoxia (Rim *vs.* Inv) (D), Inflammatory response (5ALA+ *vs.* 5ALA-) (E), and TNFA signaling via NFKB (5ALA+ *vs.* 5ALA-) (F). The heatmaps show hierarchical clustering based on z-scored expression (Log_2_ TPM+1) of the leading edge genes of Glycolysis (G). Boxplot illustrating significant expression (BH < 0.05) changes between brain regions for leading edge genes from- Glycolysis (H), Hypoxia (I), and Inflammatory response/ TNFA signaling via NFKB (J). Clustered heatmap of brain cell type (oligodendrocytes, neurons, astrocytes, and cultured astroglia) specific enrichment analysis. The color intensities depict the normalized enrichment scores (NES) of each cell type on different brain regions (Core, Rim, Inv, 5ALA-, 5ALA+) (K)

**Supplementary Figure S3.**
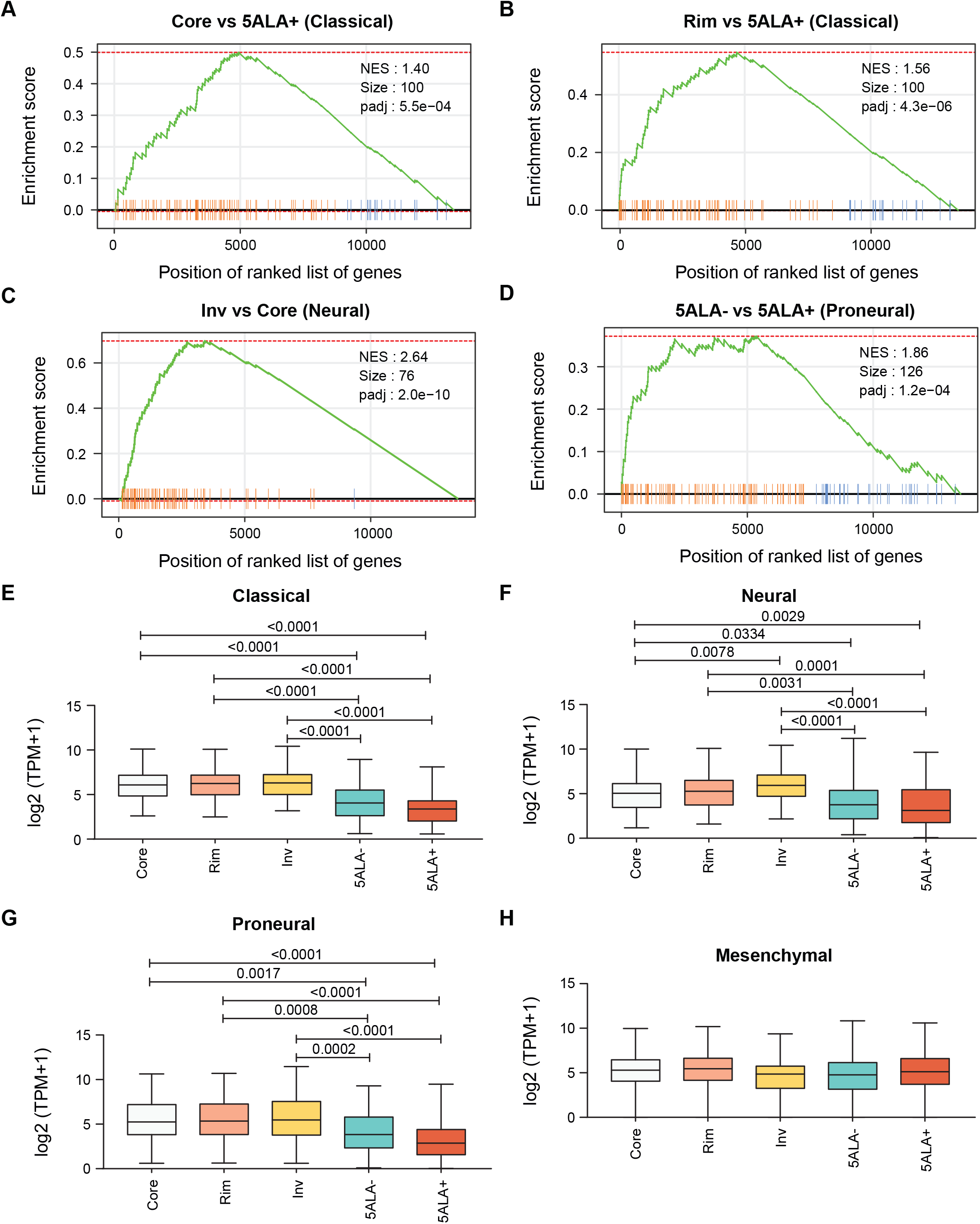
GSEA plots of gene clusters from GBM subtypes – Classical (Core *vs.* 5ALA+) (A), Classical (Rim *vs.* 5ALA+) (B), Neural (Inv *vs.* Core) (C), and Proneural (5ALA- *vs.* 5ALA+) (D). The boxplots represent the log_2_ (TPM+1) expression levels for each brain region (Core, Rim, Inv) and 5ALA sorted cells (5ALA-, 5ALA+), with significant expression differences (P-value < 0.05) denoted among GBM regions and 5ALA sorted cells for GBM subtypes – Classical (E), Neural (F), Proneural (G) and Mesenchymal (H).

**Supplementary Figure S4.**
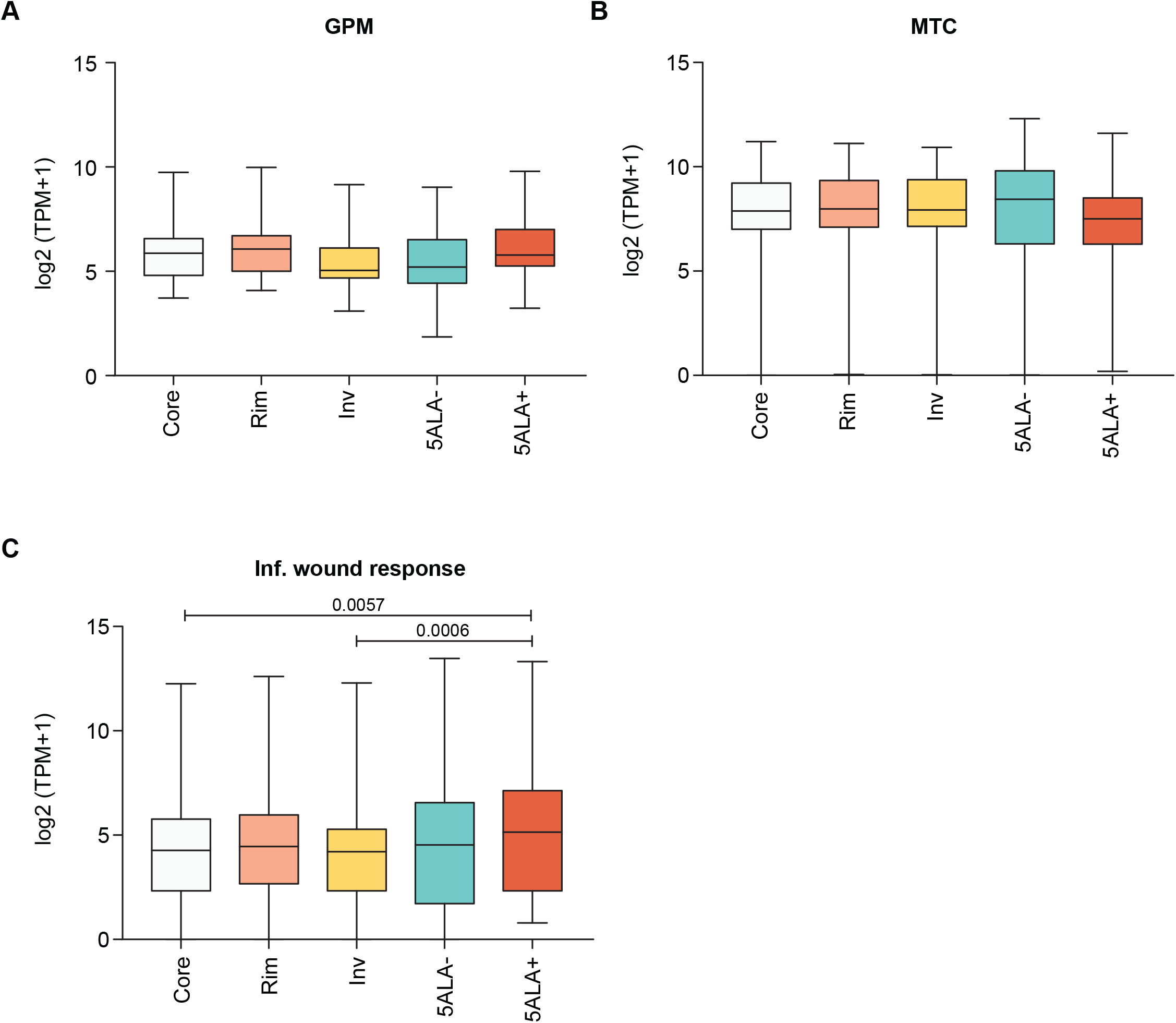
Box plots showing the relative log_2_ (TPM+1) expression levels of three cellular and metabolic state gene sets – GPM (A), MTC (B), and Inflammatory wound response (C) for Core, Rim, Inv, 5ALA- and 5ALA+ cells. Respective p-values (if <0.05) are also shown.

**Supplementary Figure S5.**
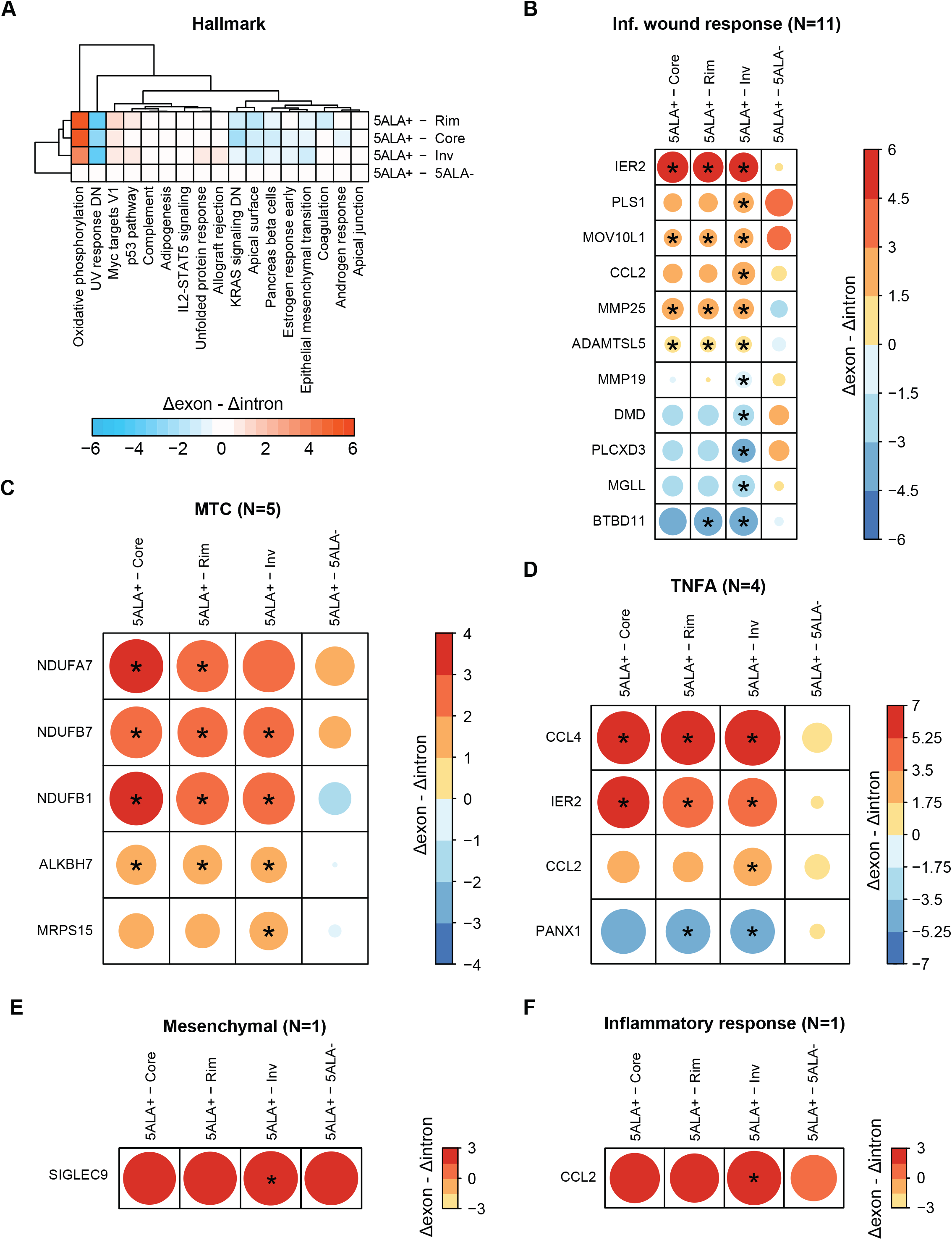
Hallmark enrichment is shown via a color code representing the difference between delta exon and intron ((Δexon-Δintron). Thus, positive (red) and negative (blue) values indicate higher exon and intron changes respectively in 5ALA+ cells (A). Regulated DEGs with a significant change in exon and intron counts are shown as heatmaps for TNFA signaling (B), Inflammatory response (C), Mesenchymal subtype (D), Inflammatory wound response and MTC (F). The color code represents the difference between delta exon and delta intron (Δexon-Δintron). The significant changes of Δexon-Δintron are shown as (*). The size of the circle is according to the adjusted –log10 p-value.

**Supplementary Figure S6.**
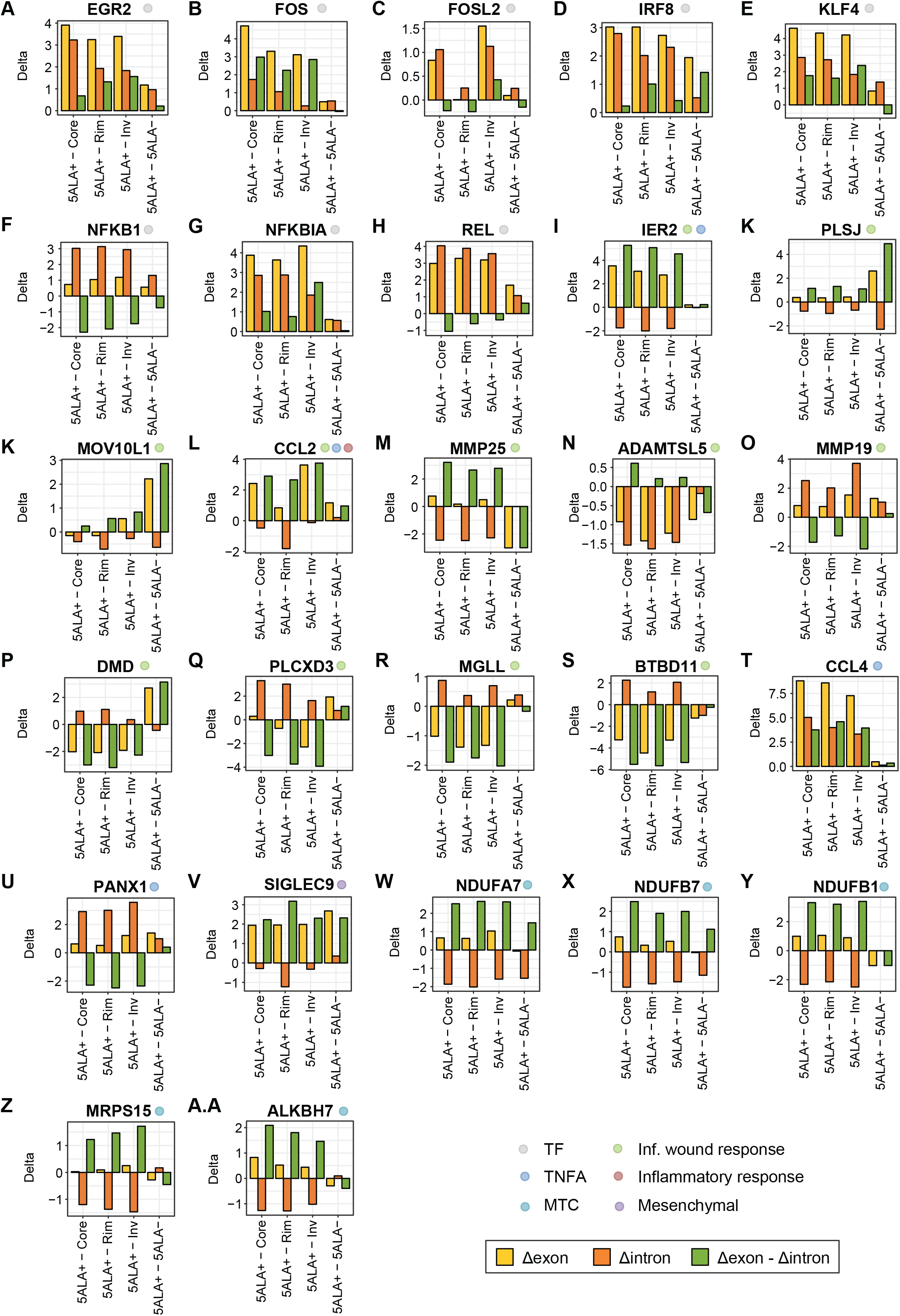
Bar plots represent delta exon (Δexon), delta intron (Δintron) and difference between Δexon and Δintron (Δexon - Δintron) for the significantly DEGs. The circles adjacent to the gene symbol represent the gene-signature and the colors of the circle indicate different gene-signatures. [TF = Transcription Factor; Inf. wound response = Infammatory Wound Response; TNFA = tumor necrosis factor – alpha; MTC = Mitochondrial Subtype].

**Supplementary Figure S7.**
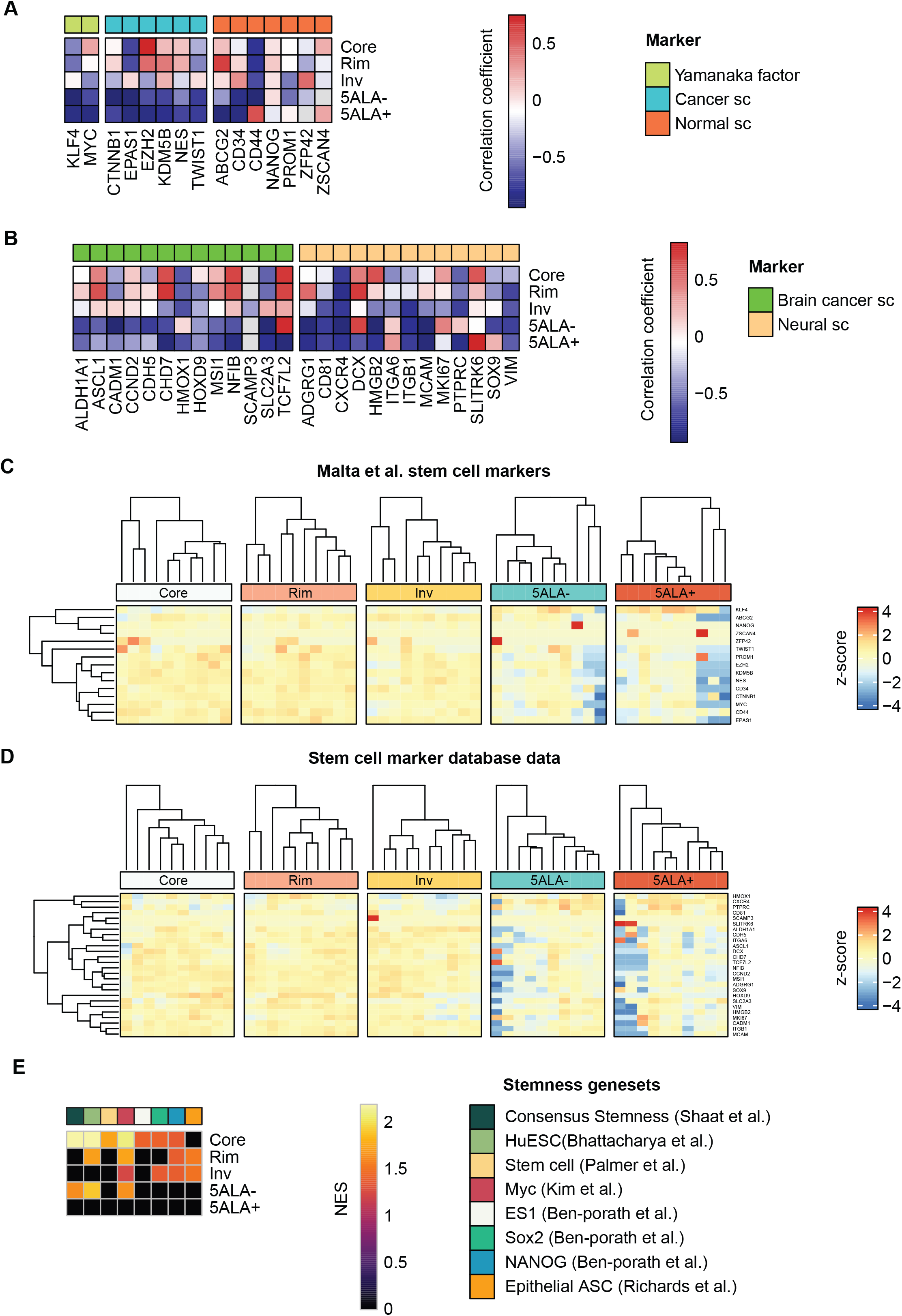
Correlation between mRNAsi values and mRNA expression for available Malta *et al.* stem cell markers (A). Correlation between mRNAsi and mRNA expression for markers from stem cell marker database (B). Heatmap of RNA-seq expression z-scores for the Malta *et al.* stem cell marker signatures (C) and markers from stem cell marker database (D). Heatmap illustrating normalized enrichment scores (NES) of GSEA with padj<0.05 for selected stemness gene-sets from different studies €.

**Supplementary Figure S8.**
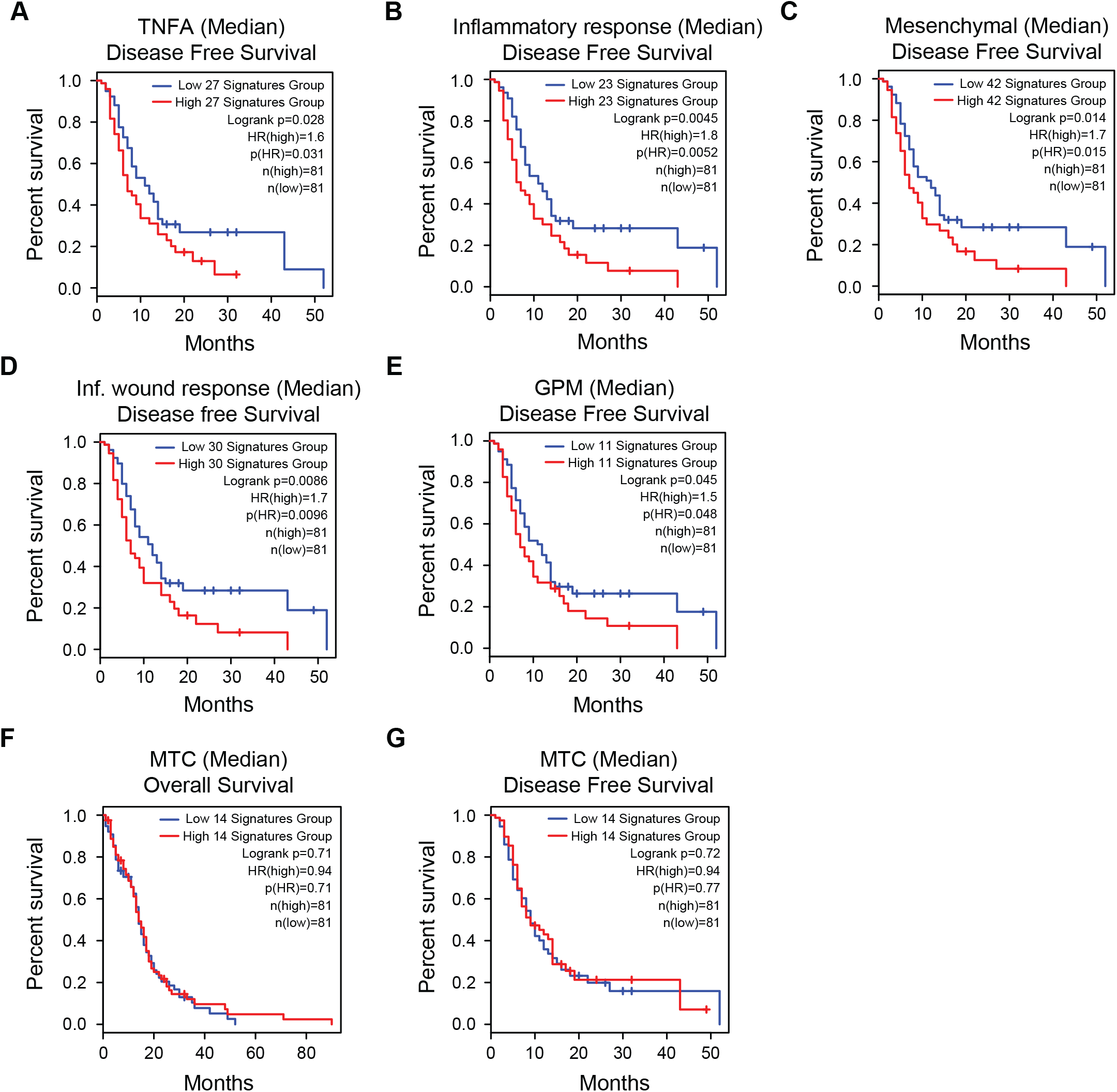
Kaplan-Meier survival analysis for disease free survival curves based on median (low expression and high expression) of five different gene signatures from TCGA-GBM samples. High TNFA leading edge signature gene expression indicated poor survival in TNFA signaling (P = 0.028) (A). High expression of Inflammatory response (P = 0.0045) (B), Mesenchymal (0.0014) (C), Inflammatory wound response (P = 0.0086) (D) and GPM (0.045) (E) gene signatures exhibited poor survival respectively.

**Supplementary Figure S9.**
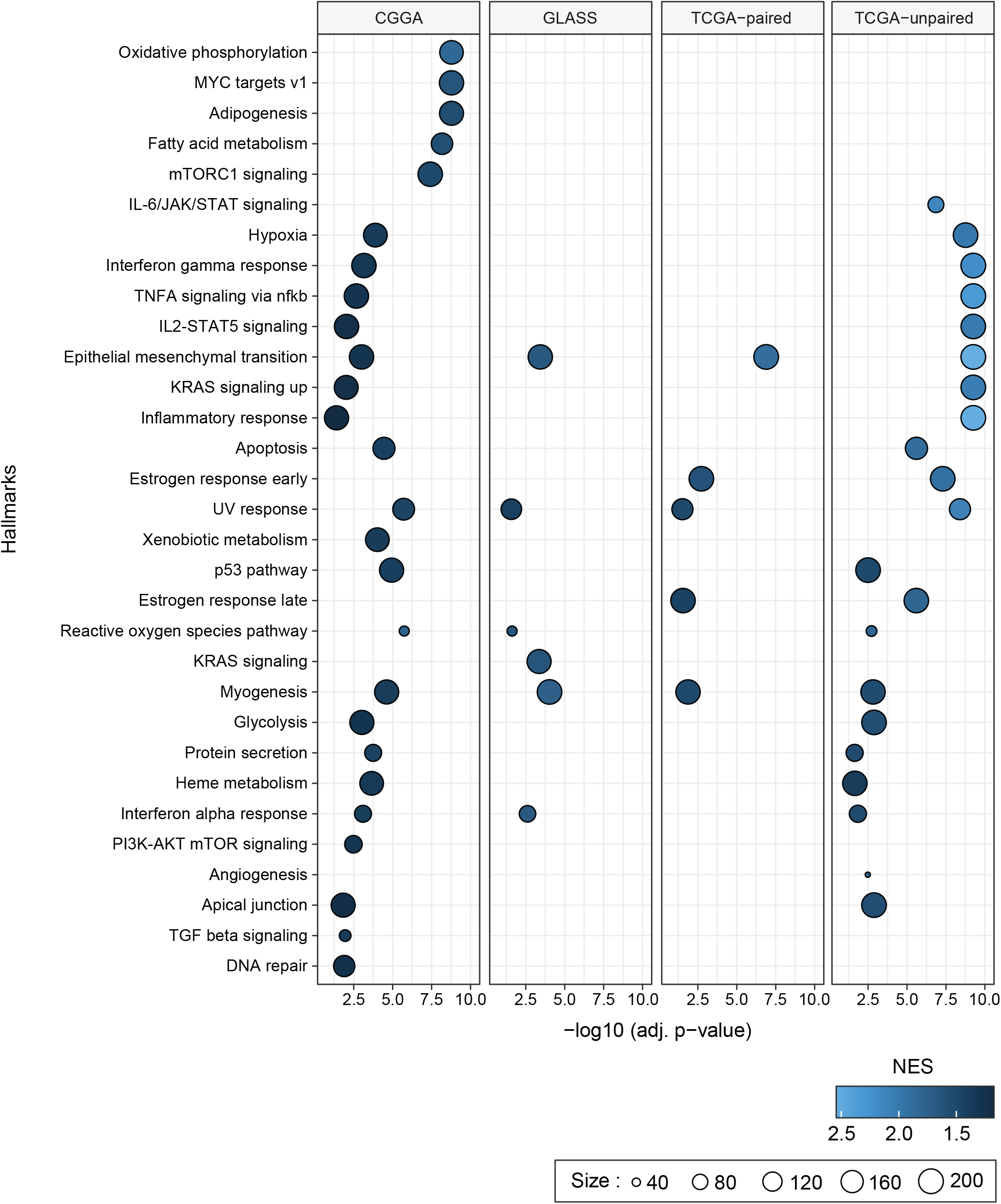
Hallmark enrichment analysis of DEGs between recurrent and primary GBM tumors. The hallmarks that are enriched in recurrent tumors in comparison to parimary tumors from TCGA, CCGA and GLASS cohorts are illustrated. Each bubble indicate one particular hallmark (y-axis). The x-axis indicate a –log10 adjusted p-value. The color gradiant of the bubble indicates the normalized enrichment score (NES) whereas the size of the bubble depends on the gene-count for each hallmark. Paired and unpaired analysis was performed for the TCGA cohort.

## Supplemnetary Tables

**Supplementary Table 1:** Compiled Gene-signatures

**Supplementary Table 2:** Hallamark GSEA and leading gene-edge gene list

**Supplementary Table 3:** Neural cell type GSEA and leading gene-edge gene list

**Supplementary Table 4:** GBM-subtype GSEA and leading gene-edge gene list **Supplementary Table 5:** Cellular and metabolic gene-set GSEA and leading gene- edge gene list

**Supplementary Table 6:** TF-target gene interaction

**Supplementary Table 7:** Enrichment of selected gene-signatures in DEGs with significantly altered exon and intron counts in 5ALA+ cells

**Supplementary Table 8:** Enrichment of stemness asscoaietd gene-signatures

**Supplementary Table 9:** Genes correlated with stemness in 5ALA+ cells

**Supplementary Table 10:** Genes asscoaited with poor survival

**Supplementary Table 11:** GSEA in primary and recurrent tumors

## Notes

### Competing Interest Statement

The authors have declared no competing interest.

